# Advancing DIA-based Limited Proteolysis Workflows: introducing DIA-LiPA

**DOI:** 10.1101/2025.11.11.687786

**Authors:** Chloé Van Leene, Emin Araftpoor, An Staes, Andrea Argentini, Marcel Bühler, Lieven Clement, Kris Gevaert

## Abstract

Limited proteolysis coupled to mass spectrometry (LiP-MS) probes protein conformational dynamics, but interpretation of LiP-MS data is complicated by heterogenous proteolytic cleavage patterns and missing data. Recent advances in data-independent acquisition (DIA) and machine learning-based search engines promise improved sensitivity and reproducibility, yet their performance in LiP-MS workflows remains underexplored. We systematically evaluated selected library-free DIA workflows using a rapamycin-treated human cell lysate and a yeast heat shock dataset, benchmarking DIA-NN and Spectronaut for identification depth, reproducibility and false discovery rate control. Our results show that library-free approaches achieve high sensitivity, eliminating the experimental overhead and sample requirements associated with empirical libraries. Building on these advances, we introduce a DIA-based Limited Proteolysis data Analysis pipeline (DIA-LiPA), a data analysis workflow tailored for LiP-MS data that integrates semi-tryptic- and tryptic-level precursor data and accounts for missingness to enable structural interpretation. Validation across multiple datasets confirmed that DIA-LiPA reproduces known structural signatures and uncovers additional regulatory patterns, providing a robust framework for mechanistic insights into protein dynamics.

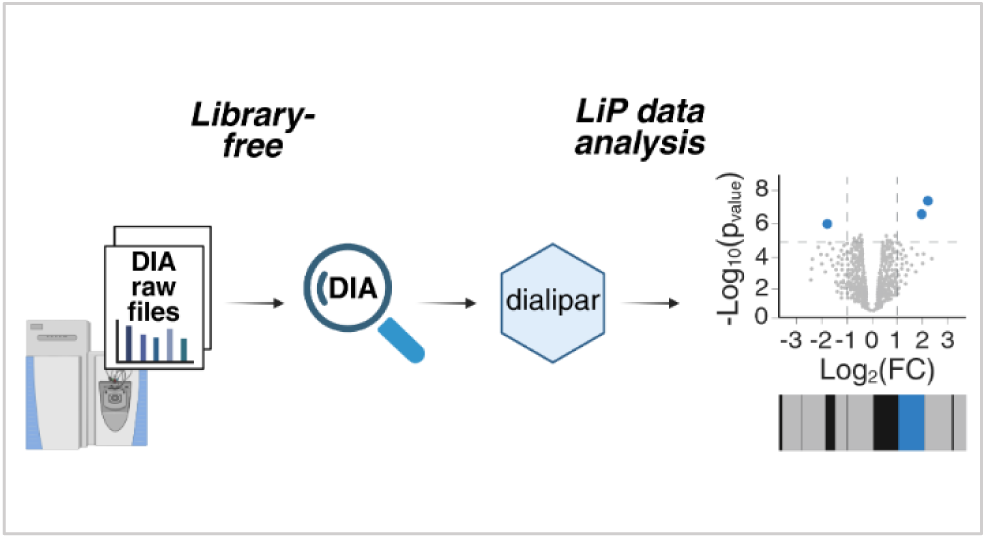

## Introduction

Protein functions are tightly determined by their structures. Instead of having a single, static form, proteins adopt various conformational states, known as conformational ensembles, that may change in response to environmental stimuli, ligand binding or post-translational modifications ^1^. Such conformational changes are strongly linked to alterations in protein functions and, for example, regulate protein-protein interactions ^2,3^. Therefore, understanding protein conformational dynamics helps elucidating cellular processes and disease mechanisms^1,4^.

Structural proteomics aims to characterize conformational changes and several mass spectrometry-based techniques have been developed for this purpose, including cross-linking mass spectrometry (XL-MS) ^5,6^, hydrogen-deuterium exchange mass spectrometry (HDX-MS) ^7^, thermal proteome profiling (TPP) ^8,9^, peptide-centric local stability assay (PELSA) ^10^ and limited proteolysis coupled to mass spectrometry (LiP-MS) ^11^. While XL-MS and HDX-MS provide valuable insights into protein-protein interactions and solvent accessibility, respectively, their proteome-wide application remains technically challenging due to sample complexity, data analysis and limited coverage of the proteins and structural regions accessible to these methods ^12^. TPP and its more recent derivative PISA (Proteome Integral Solubility Assay) measure changes in protein stability upon perturbation, though do not directly provide conformational information ^12–14^. PELSA, a more recent limited-proteolysis-based strategy uses high concentrations of trypsin to sensitively detect ligand-induced local stability shifts. This enables identification of ligand-binding regions and their corresponding proteins. While conceptually similar to LiP-MS, PELSA is less sensitive in pinpointing exact binding sites and conformational changes at the peptide level ^10^.

Among these techniques, LiP-MS stands out for its ability to probe conformational changes in complex samples. It relies on the principle that a protein’s conformation determines those sites at its surface that will be cleaved by a protease, and such sites are found in solvent-exposed and flexible protein regions. Conformational changes altering protease access to these regions thus result in distinct protease cleavage patterns. In LiP-MS, proteins are first briefly incubated with a promiscuous protease (e.g., proteinase K) under native conditions, followed by complete digestion with trypsin. This results in so-called conformotypic peptides, a term that encompasses both newly formed semi-tryptic and tryptic peptides whose abundance is affected by structural rearrangements. Conformational peptides thus serve as structural fingerprints of a protein. By now comparing proteolytic LiP signatures across conditions, such as drug or metabolite binding, LiP-MS enables the detection of conformational changes at peptide level and the identification of functionally relevant protein regions ^11,15,16^.

Initially, LiP-MS workflows employed Data-Dependent Acquisition (DDA) to identify conformotypic peptides and build spectral libraries for downstream analysis ^11,15^. DDA only selects and fragments the most abundant peptide ions, hereafter referred to as precursors, and its stochastic precursor selection leads to missing values and thus limited reproducibility in complex samples. To overcome these limitations, Data-Independent Acquisition (DIA) has been increasingly adopted. DIA provides deeper proteome coverage and markedly improved reproducibility compared to DDA because all precursors within defined m/z windows are fragmented systematically and thus independently of their signal intensity. This reduces missing values and improves consistency across runs, particularly for complex samples. DIA additionally enables the detection of low-abundance peptides that are often missed by DDA, making it highly suitable for unbiased and quantitative proteome profiling ^17–19^. These advantages have also motivated the adoption of DIA in LiP-MS workflows ^16,20^. These are often supported by project-specific spectral libraries generated from DDA runs, which provide deep coverage, especially when peptides were pre-fractionated prior to DDA ^16^. However, generating such libraries is time-consuming, requires additional sample material and remains subject to DDA’s inherent biases.

Recent developments have introduced library-free DIA approaches for LiP-MS, which rely on *in silico* predicted libraries rather than experimentally generated ones. These workflows reduce experimental overhead while maintaining comprehensive coverage. Similar advantages of using *in silico* predicted libraries over conventional DDA workflows have also been reported for other MS applications, including immunopeptidomics ^18,21^ and phosphoproteomics ^22^, further supporting the robustness of this approach. The semi-tryptic nature of LiP-MS data has posed challenges for library-free DIA approaches, given the increased spectral complexity and expanded search space ^19,20,23^. Recent advances in machine learning and deep neural networks have addressed these issues. Tools like DIA-NN ^24^ and Spectronaut ^25^ now enable accurate precursor identification and quantification directly from DIA LiP-MS data, without the need for experimental spectral libraries. These tools can predict precursor fragmentation patterns and retention times with high precision, enabling library-free workflows that match and even exceed the sensitivity and reproducibility of traditional DDA-based approaches ^23,24^. DIA-NN, uses a peptide-centric approach to identify candidate precursors, followed by a spectrum-centric strategy to resolve interference from co-eluting peptides ^24^. In contrast, Spectronaut uses a spectrum-centric approach by generating DDA-like pseudo-MS/MS spectra, which can be searched using conventional DDA search engines.

However, false discovery rate (FDR) control remains a challenge in DIA workflows, particularly in conformational proteomics applications like LiP-MS, where it is important to distinguish subtle conformational changes from background variation and analytical noise ^19,26^. In DIA, such noise can arise from the co-isolation and co-fragmentation of multiple precursors within the same selection window, leading to inference from co-eluting peptides and increased spectral complexity ^20^. This challenge is further amplified in LiP-MS by the inclusion of semi-tryptic peptides, which increase peptide-level complexity and expand the search space. As a result, FDR estimation becomes more difficult, with the search space inflating up to 20-fold compared to tryptic searches, thereby increasing the likelihood of identifying false positives ^20^. While DDA-based libraries help control the FDR by constraining the search space, library-free approaches rely on the discriminative power of deep learning to more effectively distinguish true from false positives.

To overcome the limitations of DDA-based library generation and address the specific challenges of LiP-MS data, we developed DIA-LiPA, a flexible and comprehensive pipeline tailored for DIA-based Limited Proteolysis data Analysis. DIA-LiPA operates on precursor-level quantification outputs from DIA-NN or Spectronaut and performs data filtering, abundance normalization, differential abundance (DA) analysis as well as relative differential abundance (RDA) analysis by correcting LiP precursor-level log_2_ fold changes (FC) for changes in overall protein abundance using trypsin controls, ultimately generating interpretable outputs such as volcano plots and structural barcodes with missingness information included. The pipeline, implemented in an open-source R/Bioconductor msqrob2 workflow dialipar^1^, is compatible with both Spectronaut and DIA-NN, and adaptable to various experimental designs. Despite the fact that the recent release of DIA-NN versions 2.3.0 and upwards^2^ include a semi-tryptic digestion option, DIA-LiPA was initially developed in the context of a semi-tryptic FASTA-based workaround in DIA-NN 2.2.0, hence we also incorporated this approach into our study. Notably, several pipelines or software packages have been proposed for LiP-MS data analysis, including protti ^27^, MSstatsLiP ^16^ and LiPAnalyzeR ^28^, each offering tailored approaches for peptide-level quantification and structural interpretation. However, none of these currently support DIA-NN output or report missingness patterns, which are key features addressed by DIA-LiPA.

In this study, we use DIA-LiPA as a unified downstream analysis pipeline to systematically compare traditional LiP-MS acquisition workflows, including DDA-based spectral libraries and library-free approaches in both DIA-NN (version 2.3.0) and Spectronaut (version 20.3). Our results demonstrate that in our datasets, focused on LiP-MS and semi-tryptic peptide analysis, library-free workflows perform comparably or better than DDA-based library methods. This suggests that empirical libraries may not be strictly necessary for these types of applications. Importantly, DIA-LiPA allows for the interpretation of missingness patterns, which are biologically relevant in LiP-MS. Structural changes may completely prevent the generation of specific semi-tryptic peptides, resulting in condition-specific absence that reflects conformational rearrangements rather than technical noise. Furthermore, by not restricting the analysis to fully tryptic peptides, DIA-LiPA provides a more comprehensive and robust view of conformational dynamics, capturing abundance changes, shifts in proteolytic accessibility and allowing mutual validation between peptide types.

## Materials and methods

### Cell Culture and Proteome Preparation

HEK293T cells were cultured in Dulbecco’s Modified Eagle Medium (DMEM; Gibco), supplemented with 10% fetal bovine serum (FBS; Gibco) under standard conditions (37 °C, 5% CO2). After several passages, cells were harvested by trypsinization, followed by neutralization with complete DMEM. The cell suspension was centrifuged at 450 *g* for 3 min at room temperature. The supernatant was aspirated and the pellet was washed with phosphate-buffered saline (PBS), followed by a second centrifugation under the same conditions. After removal of the supernatant, the resulting cell pellet was snap-frozen in liquid nitrogen and stored at -80 °C until further use.

A cell pellet equivalent to approximately 12 million HEK293T cells was lysed using a pellet pestle motor (Kimble^®^) in 300 µL of LiP buffer (100 mM HEPES, pH 7.4; 150 mM KCl; 1 mM MgCl_2_). Lysis was performed on ice with 10 pulses every minute for 10 min. The lysate was then centrifuged at 16000 *g* for 10 min at 4 °C, and the supernatant was transferred to a fresh microcentrifuge tube (Eppendorf, 1.5 mL). Protein concentration was determined using the Pierce BCA Protein Assay Kit (Thermo Scientific), following the manufacturer’s instructions.

### Limited Proteolysis (LiP) Treatment

The cell lysate was divided into sixteen samples: eight trypsin control (TC) samples, subjected to only trypsin digestion, and eight LiP samples, subjected to double-protease digestion with proteinase K followed by complete trypsin digestion. Each sample contained 50 µg of lysate.

The LiP protocol ^16^ was performed as follows: samples were incubated at 25 °C for 5 min, followed by treatment with either 10 µM rapamycin (Sigma-Aldrich) in 0.1% dimethyl sulfoxide (DMSO; Sigma-Aldrich) or 0.1% DMSO as vehicle control, at 25 °C for 5 min. Proteinase K (PK; Promega) was then added to the LiP samples at an enzyme-to-substrate ratio of 1:100 (w/w) and incubated at 25 °C for 5 min. For the control samples, an equivalent volume of water was added instead of PK.

Proteolysis was stopped by heating the samples at 99 °C for 5 min, followed by cooling at 4°C for 5 min. Deoxycholate (DOC; Sigma-Aldrich) was then added to a final concentration of 5%.

### Trypsin Digestion and Peptide Preparation

Samples were reduced with 5 mM tris-(2-carboxyethyl)phosphine (TCEP, Thermo Scientific) at 37 °C for 30 min, and alkylated with 40 mM iodoacetamide (Sigma-Aldrich) at 30 °C for 30 min in the dark. Samples were diluted with 0.1 M ammonium bicarbonate (Merck) to reduce the DOC concentration to 1%.

Sequencing Grade Modified Trypsin (Promega) was added at an enzyme-to-substrate ratio of 1:100 (w/w) and incubated overnight at 37 °C in a thermomixer at 800 rpm. Digestion was stopped by acidifying the samples to pH < 2 with 50% formic acid.

Peptides were desalted using Microspin C18 columns (Nest Group), eluted with 50% acetonitrile and 0.1% formic acid, dried in a vacuum centrifuge and stored at -20 °C.

### High-pH Reversed-Phase Fractionation of Peptides

Peptides were reconstituted in 20 µL loading solvent A (0.1% trifluoroacetic acid in water/acetonitrile (ACN) (98:2, v/v)). LiP and control replicates (16 µL per sample, 4 replicates per condition) were pooled separately, yielding approximately 50 µg of total peptide digest per condition. Each pooled sample was dried down, reconstituted in fractionation solvent A (10 mM ammonium acetate pH 5.5 in H_2_O) and fractionated into 12 pooled fractions using high-pH reversed-phase chromatography on a Vanquish™ Flex (Thermo Fisher Scientific) equipped with a Zorbax 300SB-C18 column (0.3 × 150 mm, 3.5 µm; Agilent) preceded by a Zorbax 300SB-C18 guard cartridge (0.5 x 5 mm, 5 µm; Agilent).

Separation was performed using a 100-minute linear gradient from 1 to 100% acetonitrile (ACN) at a flow rate of 8 µL/min. Peptide elution was monitored with 1-min fractions collected from 50 to 148 min and online pooled every 12 min. After vacuum drying, the individual fractions were resuspended in loading solvent A’ (0.5% ACN in 0.1% trifluoroacetic acid (TFA)), spiked with Biognosys’ iRT kit peptides according to the manufacturer’s instructions.

### Online Liquid Chromatography

Peptides were reconstituted in 20 µL loading solvent A (0.1% trifluoroacetic acid in water/acetonitrile (ACN) (98:2, v/v)). For each injection, either 2 µL of unfractionated, unpooled sample (for DIA) or 10 µL of fractionated, pooled sample (for DDA) was injected on an Ultimate 3000 Pro Flow nanoLC system in-line connected to a Q Exactive HF mass spectrometer (Thermo). Trapping was performed at 20 μL/min for 2 min in loading solvent A on a 5 mm trapping column (Thermo scientific, 300 μm internal diameter (i.d.), 5 μm beads). The peptides were separated on a 250 mm Aurora Ultimate analytical column (1.7 µm C18, 75 µm i.d.; Ionopticks) kept at a constant temperature of 45 °C in a butterfly oven (Phoenix S&T). Peptides were eluted by a non-linear gradient starting at 0.5% MS solvent B (0.1% FA in acetonitrile) reaching 26% MS solvent B in 75 min, 44% MS solvent B in 95 min, 56% MS solvent B in 100 min followed by a 5-minute wash at 56% MS solvent B and re-equilibration with MS solvent A (0.1% FA in water) at a constant flowrate of 250 nL/min.

### DIA Acquisition

Unfractionated, unpooled peptides were analysed in data-independent acquisition (DIA) mode, automatically switching between MS and MS/MS acquisition. Full-scan MS spectra ranging from 375-1,500 m/z with a target value of 5E6, a maximum fill time of 50 ms and a resolution of 60,000 were followed by 30 quadrupole isolations with a precursor isolation width of 10 m/z for HCD fragmentation at an NCE of 30% after filling the trap at a target value of 3E6 for maximum injection time of 45 ms. MS2 spectra were acquired at a resolution of 15,000 at 200 m/z in the Orbitrap analyser without multiplexing. The isolation intervals ranging from 400-900 m/z of 10 m/z were created with the Skyline software tool. QCloud was used to control instrument longitudinal performance during the project ^29,30^.

### DDA Acquisition for Spectral Library Generation

Pooled, fractionated peptides were analysed in data-dependent acquisition (DDA) mode, automatically switching between MS and MS/MS acquisition for the 12 most abundant ion peaks per MS spectrum. Full-scan MS spectra (375-1,500 m/z) were acquired at a resolution of 60,000 in the Orbitrap analyzer after accumulation to a target value of 3E6 with a maximum ion time of 60 ms. The 12 most intense ions above a threshold value of 1.3E4 were isolated (isolation window of 1.5 m/z) for fragmentation at a normalized collision energy of 30% after filling the trap at a target value of 1E5 for maximum 80 ms. MS/MS spectra (with a fixed first mass of 145 m/z) were acquired at a resolution of 15,000 in the Orbitrap analyzer. The polydimethylcyclosiloxane background ion at 445.120028 Da was used for internal calibration (lock mass) and QCloud has been used to control instrument longitudinal performance during the project ^29,30^.

### Mass Spectrometry Data Analysis

DIA raw files were searched in two different ways: (1) library-free using *in silico* predicted spectral libraries, and (2) empirical library-based using spectral libraries generated from DDA analysis of offline peptide fractions.

#### Library-free (in silico predicted) DIA

DIA data were analysed using Spectronaut (version 20.3; Biognosys AG) ^25^ and DIA-NN (version 2.3.0) ^24^ (**Table 1**).

**Table 1:** Overview of different data analysis tools and settings used in this study.

|  | Tool | Version | Digestion setting | Library size (if applicable) | Notes |
| --- | --- | --- | --- | --- | --- |
| *In silico* predicted library | Spectronaut | 20.3 | Semi-specific (full enumeration) | / |  |
|  | DIA-NN | 2.3.0 | Semi | 51,154,169 precursors | Semi-option newly available in this version. |
|  | DIA-NN | 2.2.0 | / | 47,923,754 precursors | Library generated from semi-tryptic FASTA; semi-specific digestion not yet supported. |
| Empirical library generation | Spectronaut Pulsar | 20.3 | Semi-specific (LiP) / Specific (TC) | / | Libraries used in Spectronaut. |
|  | MSFragger<br>(FragPipe) | 4.3<br>(23) | SEMI (LiP) /<br>ENZYMATICAL<br>(TC) | / | DIA-SpecLib-Quant workflow;<br>libraries used in<br>DIA-NN. |
| Empirical library<br>search | Spectronaut | 20.3 | Semi-specific<br>(LiP) / Specific<br>(TC) | 47,044 (LiP) /<br>51,377 (TC)<br>precursors |  |
|  | DIA-NN | 2.3.0 | Semi | 49,301 (LiP) /<br>48,823 (TC)<br>precursors |  |

In both tools, semi-specific digestion was applied to enable the identification of semi-tryptic peptides directly from DIA data. Notably, semi-specific digestion was only recently introduced in DIA-NN (version 2.3.0 and upwards); earlier versions required a semi-tryptic FASTA workaround (see Supporting Information). For DIA-NN, the scoring strategy was set to Proteoforms. Both tools used the human UniProt FASTA (01/2024) supplemented with the proteinase K (*Tritirachium album*) sequence. Search parameters were harmonized across tools and included missed cleavages 1, charge states 1 to 4, peptide length 7 to 30 amino acids, fragment ion m/z range 200 to 2,000, maximum number of variable modifications 1, oxidation (M) as variable modification, carbamidomethylation (C) as fixed modification and N-terminal methionine excision disabled.

All other settings were kept at default values, except for two modifications made specifically in Spectronaut 20.3. First, the Semi-Specific Pipeline was set to Full Enumeration (SN19) rather than the default Smart Enumeration (SN20), because Smart Enumeration drastically suppressed semi-tryptic identifications when processing raw files (down to 5%). Full Enumeration restored semi-tryptic identifications to 35%. Second, applying the HTRMS converter prior to import further increased semi-tryptic identifications to 38%. The HTRMS conversion was required for our in-house generated raw files containing negative polarity scans in the first few minutes of the MS acquisition, which otherwise interfere with Spectronaut’s raw-file parsing and lead to reduced identifications (Supplementary Table S1).

To further demonstrate the applicability and usefulness of the DIA-LiPA pipeline, we reanalysed a yeast heat shock dataset (PRIDE accession: PXD022297) ^31^ using library-free DIA in both Spectronaut and DIA-NN. To ensure comparability with the original publication, we employed the same *S. cerevisiae* (strain S288c) UniProt FASTA (11/2016).

#### Empirical library-based DIA

Offline DDA fractionation data were analysed using either Spectronaut Pulsar (version 20.3) or FragPipe (version 23) using MSFragger version 4.3 ^32^ for spectral library generation (Table 1). To ensure a clean comparison with the library-free DIA searches, the same search parameters were applied as described above. These libraries were then used to search the DIA data using Spectronaut and DIA-NN (version 2.3.0) (Table 1).

In Spectronaut, spectral libraries were generated using Pulsar with the same UniProt FASTA (supplemented with the proteinase K sequence). LiP samples were searched using semi-specific digestion and trypsin-control samples with specific digestion.

In FragPipe, the DIA-SpecLib-Quant workflow was used to build spectral libraries from DDA files. The same FASTA file was used as in Spectronaut supplemented with the Biognosys iRT peptide sequences, with cleavage set to SEMI for LiP samples and ENZYMATIC for trypsin-control samples. In the Spec Lib tab, RT calibration was set to Biognosys_iRT and psm.tsv was selected as the Filetype to Convert.

### Entrapment to Assess False Discovery Rate (FDR)

To assess FDR control in the library-free approaches, the *E. coli* proteome (08/2025) was included as an entrapment database alongside the previously mentioned human proteome database. Because the entrapment database is smaller than the human proteome database, the combined estimation method was applied, which accounts for the ratio *r* of entrapment to target database sizes ^26^. The estimated false discovery proportion (FDP) per sample was calculated as:

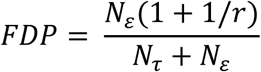

where *N_τ_* and *N_ε_* denote the number of precursors identified from the human and entrapment (*E. coli*) databases, respectively. Subsequently, the FDPs were averaged over all samples to get an estimate of the FDR, which is the expected false discovery proportion.

For entrapment validation, all protein-level filters were disabled: in Spectronaut by setting Protein Qvalue Cutoff (Experiment and Run), Protein PEP Cutoff and Protein Group FDR to 1.0; in DIA-NN by performing the search in Proteoforms scoring mode ^33^ with the --no-refine-q option to avoid protein-level refinement during validation. Precursor-level FDR thresholds remained at 1%.

### DIA-LiPA Statistical Method Description

Precursors with a valid MS2-based quantity (Precursor.Quantity > 4 in DIA-NN or FG_MS2RawQuantity > 4 in Spectronaut) are filtered to retain those with at least two identifications per sample group. For example, a precursor with three measurements in group

A and one in group B, will only be filtered out of group B. Precursors that can be the product of tryptic digestion are annotated as tryptic, those that either end on or are preceded by lysine or arginine as semi-tryptic, and others as non-tryptic. Precursors that are not proteotypic or repeat within a protein sequence are annotated within each sequence context separately. Within each pipeline, i.e., limited proteolysis (LiP) or trypsin-control (TC), data are normalised separately by subtracting a sample-specific scaling factor from all log_2_ precursor quantities. This scaling factor is calculated using only the precursors that are shared between all samples in a pipeline (i.e., complete cases) and equals the log_2_ difference between the sample median and the pipeline-wide median. This strategy was chosen over global median normalization to avoid bias introduced by differences in precursors that are picked up across samples. By using only shared precursors across all samples, we ensure balanced normalization across peptide types and improve quantitative accuracy. Following normalisation, coverages are calculated on a protein accession level. Sample correlations are calculated based on the complete precursors using the Spearman correlation, and multidimensional scaling plots are based on the Euclidian distances of the log_2_ transformed and normalised precursors intensities that were available in the sample pairs involved in each distance. Differential abundance analysis on the LiP precursors is based on methodology from the post-translational modification (PTM) literature. Similar to Demeulemeester *et al.* ^34^ we make a clear distinction between directly assessing differential abundance (DA) at LiP precursor level and relative differential abundance (RDA) analysis so as to correct the log_2_ fold change (FC) at LiP precursor-level for overall protein-level abundance changes. The DA analysis consists of directly inferring log_2_ fold changes (log_2_ FC) on the precursor level LiP data. The LiP RDA analysis, however, depends on the design. On the one hand, when LiP and TC samples are unpaired, the strategy developed by MSstatsTMT ^35^ is used, where log_2_ FC and their corresponding standard errors (SE) are estimated both at precursor-level using the LiP data as well as upon protein-level aggregation using the TC data. Subsequently, the RDA is estimated by subtracting the corresponding TC protein-level log_2_ FC from the precursor level LiP log_2_ FC, with corresponding standard error 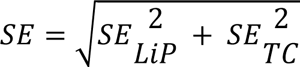 . RDA is then further inferred using t-tests, with degrees of freedom (df) based on the Satterthwaite approximation as in MSstatsTMT ^35^ or by using the more conservative dfs from the conventional LiP DA analysis. Both dfs are implemented in our msqrob2 dialipar workflow and the latter is used for all our analyses. On the other hand, for paired LiP and TC samples the msqrob2TMT approach ^34^ is adopted. Particularly, relative log_2_ LiP abundances are first obtained by subtracting the corresponding summarized log_2_ protein abundances in the paired TC samples from the precursor level log_2_ LiP intensities. Subsequently the LiP precursor-level RDA analysis proceeds similar to a conventional precursor-level DA analysis, but on the relative LiP abundances. Note, that these two approaches explicitly account for the uncertainty in the TC protein abundance quantification. The statistical modeling for both unpaired and paired TCs and LiP samples are implemented using msqrob2. Both implementations require protein group-level abundances derived from the TC samples by aggregating complete precursors to protein groups, which is done using robust summarisation on the normalised log_2_ TC precursor quantities. A schematic overview of these steps is provided in the dashed box in **Figure 1**.

All steps above are implemented in our dialipar^3^ msqrob2 workflows. The generated html reports includes several QC plots, and for each contrast a volcano plot of the differential analysis with a barcode for the proteins of interest.

The raw files, search results, FASTA files and all data analysis scripts and outputs underlying this study have been deposited to MassIVE (MSV000099740) and will be made publicly available upon acceptance of the manuscript. For peer review purposes, temporary access credentials are: username = MSV000099740_reviewer, password = bvs3Nut42YmM2d5n.

## Results and Discussion

### Experimental design for performance assessment in DIA-based LiP-MS

To evaluate the performance of our newly developed pipeline, DIA-LiPA, we used a target-engagement LiP-MS experiment in which cell lysates were treated with either rapamycin or a solvent control. Rapamycin is a well-characterized small molecule known to selectively bind FKBP1A ^36^, making it an ideal molecule to assess the sensitivity and specificity of different LiP-MS workflows. LiP-MS was performed following established protocols ^37,38^, and the resulting data were analysed using four DIA workflows: (i) empirical library search in Spectronaut or (ii) in DIA-NN, and (iii) library-free search in Spectronaut (“directDIA”) or (iv) in DIA-NN. Downstream analysis of all four workflows was performed using unpaired DIA-LiPA (**Figure 1**). For the first two workflows, we fractionated peptides using high-pH reverse-phase chromatography, followed by DDA acquisition of the resulting fractions. The acquired DDA data were then searched using either Spectronaut Pulsar or FragPipe, and the resulting identifications were used to construct a project-specific spectral library for subsequent DIA analysis. This approach integrates peptide fragmentation and retention time data, thereby restricting DIA searches to a predefined precursor list. These libraries were then used to search the DIA data using either Spectronaut or DIA-NN (**Figure 1**, grey). Peptide fractionation prior to DDA acquisition is a common practice to enhance spectral library coverage. While not explicitly included in the original LiP-MS protocol ^16^, this step substantially increases the number of identifiable precursors, resulting in a 3.5 to 4 times larger spectral library compared to the unfractionated counterpart.

In contrast, the library-free approach bypasses the need for prior DDA runs by predicting spectral libraries *in silico* from a protein FASTA file. This allows for less biased analysis of the DIA data. We implemented this strategy using Spectronaut’s directDIA mode and DIA-NN’s newly introduced semi-specific digestion feature (**Figure 1**, blue). Other LiP-MS analysis packages, such as LiPAnalyzer and MSstatsLiP, were considered but not included in our benchmarking due to incompatibility with DIA-NN outputs and data processing steps that are less accessible and more convoluted. Such constraints prevent a fair comparison under the conditions evaluated in this study.

**Figure 1:**
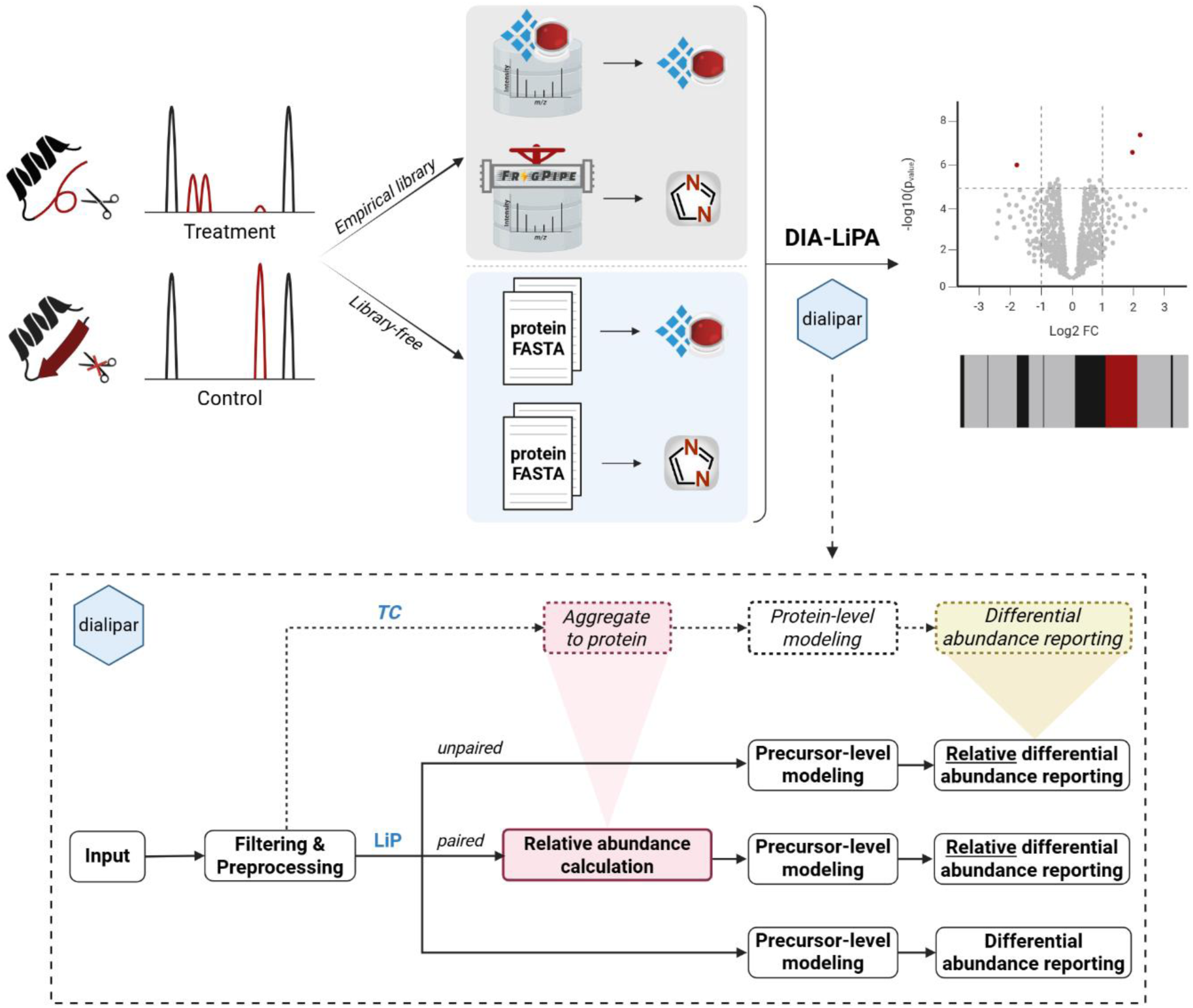
Experimental setup and DIA LiP-MS workflows analysed with DIA-LiPA. A treatment-control LiP-MS experiment was performed using cell lysates treated with either rapamycin or solvent control. The resulting data were analysed using four DIA LiP-MS workflows: grey indicates workflows using empirical DDA-based libraries, blue indicates library-free workflows using *in silico* predicted libraries. Downstream analysis was performed with unpaired DIA-LiPA to identify significant conformational changes across the proteome, with its stepwise workflow outlined in the dashed box and implemented in the *dialipar* msqrob2 workflow. The scissors icon represents limited proteolysis with proteinase K. Figure created with BioRender.com. LiP-MS; limited proteolysis coupled to mass spectrometry, TC; trypsin control, DDA; data-dependent acquisition, DIA; data-independent acquisition.

### Precursor-level identifications, coverage and reproducibility across DIA workflows in LiP-MS

We first assessed the number of identified precursors and the proportion of semi-tryptics across the different DIA workflows (**Figure 2A**). Library-free workflows identified approximately three times more precursors than empirical library-based workflows, while maintaining the expected proportion of semi-tryptics (25-30%) typical for LiP-MS experiments. Although peptide fractionation was performed to mitigate DDA stochasticity and enhance library depth, identification rates remained lower than those achieved with library-free DIA. Within the library-free approaches, Spectronaut showed slightly higher sensitivity for semi-tryptics (30%) compared to DIA-NN (28%). Furthermore, we observed a substantial overlap between library-free workflows, and about 84% of the precursors identified in empirical library-based workflows were also found in library-free workflows. This suggests that a considerable number of precursors are shared, yet many remain unique to the library-free approaches. These additional precursors were generally not of lower quality as judged by their q-values compared to those which were found by both libraries (**Supplementary Figure S1A**), nor were they of lower intensity (**Supplementary Figure S1B**). Given that library-free workflows identified three times more precursors overall, they offer broader coverage of the proteome. Importantly, these additional precursors do not proportionally increase the number of protein groups but rather provide more peptides per protein, which translates into improved sequence coverage. This is a key advantage for LiP-MS, as broader coverage enhances structural resolution and the detection of more subtle conformational changes.

To further evaluate proteome coverage, we compared the sequence coverage of shared proteins (**Figure 2B**). Library-free workflows achieved broader and more consistent coverage, due to the detection of low-abundant and semi-tryptic peptides that are often missed in empirical DDA-based libraries. As shown in the overlap panels, the choice of search engine (Spectronaut versus DIA-NN) did not appear to have a major impact on overall coverage, suggesting that differences are primarily driven by the type of library used. Similar trends were observed for non-shared proteins (**Supplementary Figure S2A**), with library-free approaches reaching higher coverages.

Semi-tryptic searches expand the search space over 15-fold compared to fully tryptic searches ^39,40^, which increases the identification rates but also introduces more false positives ^41^. To confirm that the increased sensitivity of the library-free workflows reflects true identifications, we performed an entrapment experiment by adding the *E. coli* proteome to the human protein sequence database for a semi-tryptic search. Entrapment-based FDR validation was conducted at the precursor level, with all protein-level filters disabled. The FDR was estimated using the combined entrapment method ^26^, which accounts for the database size differences and provides a conservative upper bound on the true FDR rather than a direct equivalent of the target-decoy estimate. In our semi-tryptic DIA search, the resulting precursor-level entrapment FDRs were 0.58% for DIA-NN and 1.64% for Spectronaut. These results indicate that Spectronaut’s FDR estimates might be too liberal under our semi-tryptic search conditions. However, we refrain from generalizing this observation and therefore consistently apply the default nominal 1% FDR setting for all comparative analyses. Furthermore, these values are in line with findings from a recent systematic evaluation of DIA FDR control, which shows that entrapment-based estimates typically exceeded the nominal 1% threshold ^26^. We therefore interpret these values as indicative of valid FDR control, especially given the substantially enlarged semi-tryptic search space. Together, these findings demonstrate that the library-free workflows maintain the FDR within permissible boundaries for DIA while providing substantially increased sensitivity and coverage in LiP-MS experiments, reinforcing their applicability in structural proteomics and aligning with recent benchmarking studies ^42^.

In LiP-MS, semi-tryptics arise from non-canonical cleavage events by PK, which will lower the intensities of the corresponding tryptic peptides and generate new, less or at most equally abundant, semi-tryptic peptides ^43^. Together with the fact that this workflow involves additional sample-handling steps compared to a typical proteomics sample preparation, this may lead to a reduction in quantitative precision. These factors explain why semi-tryptic workflows often exhibit broader CV distributions. Therefore, we also assessed quantitative precision by comparing precursor-level coefficients of variation (CVs) of shared precursors across workflows (**Figure 2C**), showing that CV distributions were similar across workflows. All workflows achieved median CVs below 20%, generally considered acceptable for quantitative proteomics ^44^. Similar observations were made for non-shared precursors (Supplementary **Figure S2B**), which showed higher CVs overall, with DIA-NN in library-free mode showing the smallest increase. To avoid bias introduced by global median normalization, caused by different precursors that are quantified across samples, we applied normalization based on shared precursors between conditions. This approach ensures balanced adjustment across peptide types and improves quantitative accuracy. Achieving CVs lower than 20% and maintaining FDR control is therefore particularly noteworthy, as it underscores the reliability of library-free DIA workflows, which combine high sensitivity with robust quantification while eliminating the experimental overhead of empirical library generation.

**Figure 2:**
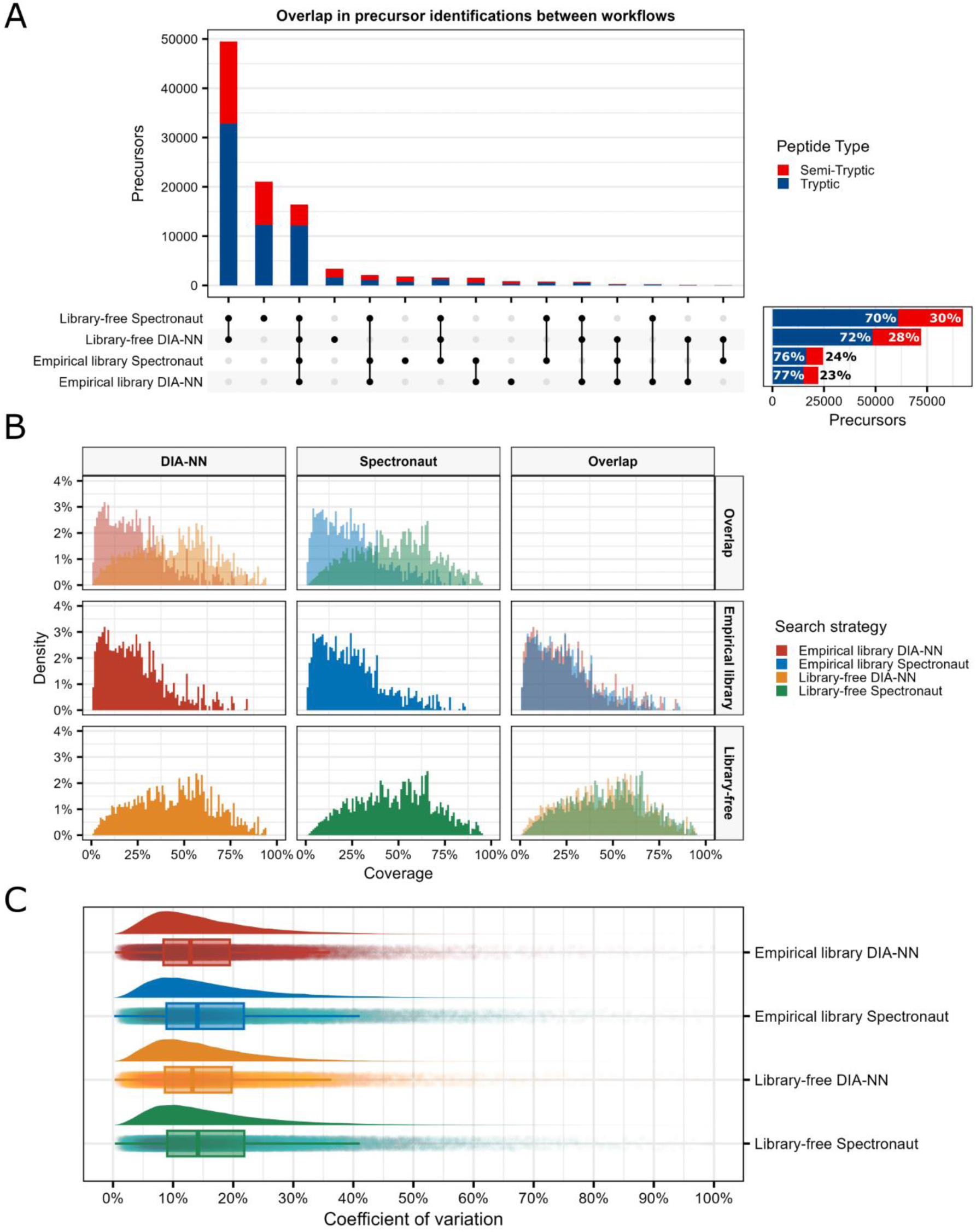
Comparison of identification, coverage and quantification precision across DIA workflows. **A)** UpSet plot showing the absolute number of precursors identified in LiP samples by each workflow and their overlaps. Percentages of tryptic and semi-tryptic precursors are indicated for each workflow. Only post-filtered data are shown (q-value < 0.01, each precursor was identified at least twice in at least one condition). **B)** Protein sequence coverage distribution for shared protein accessions in LiP samples across workflows and their overlaps. **C)** Raincloud plot showing the distribution of normalised precursor-level CVs for each workflow. The density curve represents the CV distribution, while the boxplot below indicates the median (line), interquartile range (box) and whiskers spreading to 1.5x the interquartile range. Individual points represent precursor-level CVs. CVs were calculated across all LiP samples per condition (rapamycin or vehicle control) for precursors quantified in at least three samples. CV; coefficient of variation.

### Comparison of FASTA-based and native semi-specific digestion strategies in DIA-NN for LiP-MS

We initially developed a workaround to enable semi-tryptic data analysis in DIA-NN (v2.2.0) by generating a semi-tryptic peptide FASTA file, prior to the implementation of the semi-specific digestion support in DIA-NN versions 2.3.0 and upwards. Although this workaround is no longer necessary for the latest version of DIA-NN, it remains a useful strategy for specialized applications, such as custom protease motifs (e.g., caspase motifs ^45^) or proteases not supported by current search engines, as a fully semi-tryptic search would confer an unnecessary expansion of the search space. Furthermore, the full semi-tryptic FASTA file still enables semi-tryptic searches in earlier versions of DIA-NN. We also benchmarked this workaround against the new semi-specific digestion option in DIA-NN (v2.3.0) (**Supplementary Figure S3**). Substantial overlap was observed between workflows, and the FASTA-based approach in version 2.2.0 achieved identification rates close to those obtained with the semi-specific option in version 2.3.0. This confirms that the workaround remains valid, although the new semi-specific digestion feature outperforms it and is recommended for future LiP-MS analyses.

### Capturing conformational dynamics with DIA-LiPA

We next examined the ability of each workflow to detect conformational changes using volcano plots and structural barcodes, the functional output of DIA-LiPA (**Figure 3**). All workflows identified FKBP1A with multiple relevant, i.e. significant precursors (adjusted p-value ≤ 0.05) with a relative log_2_ fold change |log_2_(FC_LiP_/FC_TC_)| ≥ 1, precursor hits, confirming the known target of rapamycin. Representative spectra for both tryptic and semi-tryptic FKBP1A precursors detected by library-free workflows are provided in **Supplementary Figure S4**.

Interestingly, precursors that were significantly upregulated in the rapamycin-treated LiP condition upon correction for TC were tryptic, whereas those that were downregulated in LiP upon correction for TC were semi-tryptic, suggesting a more shielded FKBP1A conformation upon rapamycin binding. Both library-free and empirical DIA-NN detected one additional precursor from another protein, each with low protein coverage and mapped to short, sparsely covered regions (data not shown), which may point to false positives. Importantly, although FKBP1A was consistently identified as the target of rapamycin across all search strategies, the library-free DIA-NN and Spectronaut workflows yielded substantially greater precursor coverage (30 and 34 precursors, respectively) relative to the empirical approaches (11 and 15). The expanded precursor set also contained substantially more relevant precursors (14 and 16 versus 3 in each empirical workflow), thereby improving structural resolution and enabling the detection of more subtle conformational changes.

Structural barcodes below each volcano plot visualize differential, unchanged and missing peptides across the protein sequence. Barcodes are generated by stacking precursors in the following order (bottom to top, as shown in the Relevance legend): Not Relevant, DMSO Missing, Rapamycin Missing, DMSO Up, and Rapamycin Up. Both DIA-NN and Spectronaut achieved broader FKBP1A coverage in the library-free compared to the empirical approach (88% versus 83%), enabling more distinct structural signatures. While the extra 5% coverage in library-free workflows seems minor on the stacked view of the barcode, zooming into precursor trypticity in the extended barcodes reveal additional dynamics (**Figure 4 and S5**). For example, around Trp59, a key residue in the rapamycin-binding pocket ^46^, a tryptic peptide was found at higher levels in the rapamycin condition, indicating shielding from PK cleavage. Semi-tryptic counterparts of this region are found at higher levels in the solvent condition, consistent with increased flexibility in the unbound state. Similarly, near Phe99, the tryptic peptide was not detected, but its semi-tryptic counterpart was found at higher levels in the unbound condition. These patterns provide compelling evidence of rapamycin binding, causing shielding and altered accessibility in these regions, and highlight DIA-LiPA’s ability to capture conformational changes.

Including missingness in the structural barcodes adds interpretative value when classical quantitative comparison based on relative fold-change and significance testing is not possible. While statistical modeling of missingness is challenging, missing peptides are usually filtered out downstream or imputed ^15,16^. Since imputation risks introducing bias and false positives ^47^, simply reporting absence can add interpretation value. Together, these results underscore the added value of semi-tryptics for probing conformational changes and demonstrate how DIA-LiPA integrates trypticity and missingness to deliver a more nuanced view of protein structural dynamics in DIA-based LiP-MS workflows.

**Figure 3:**
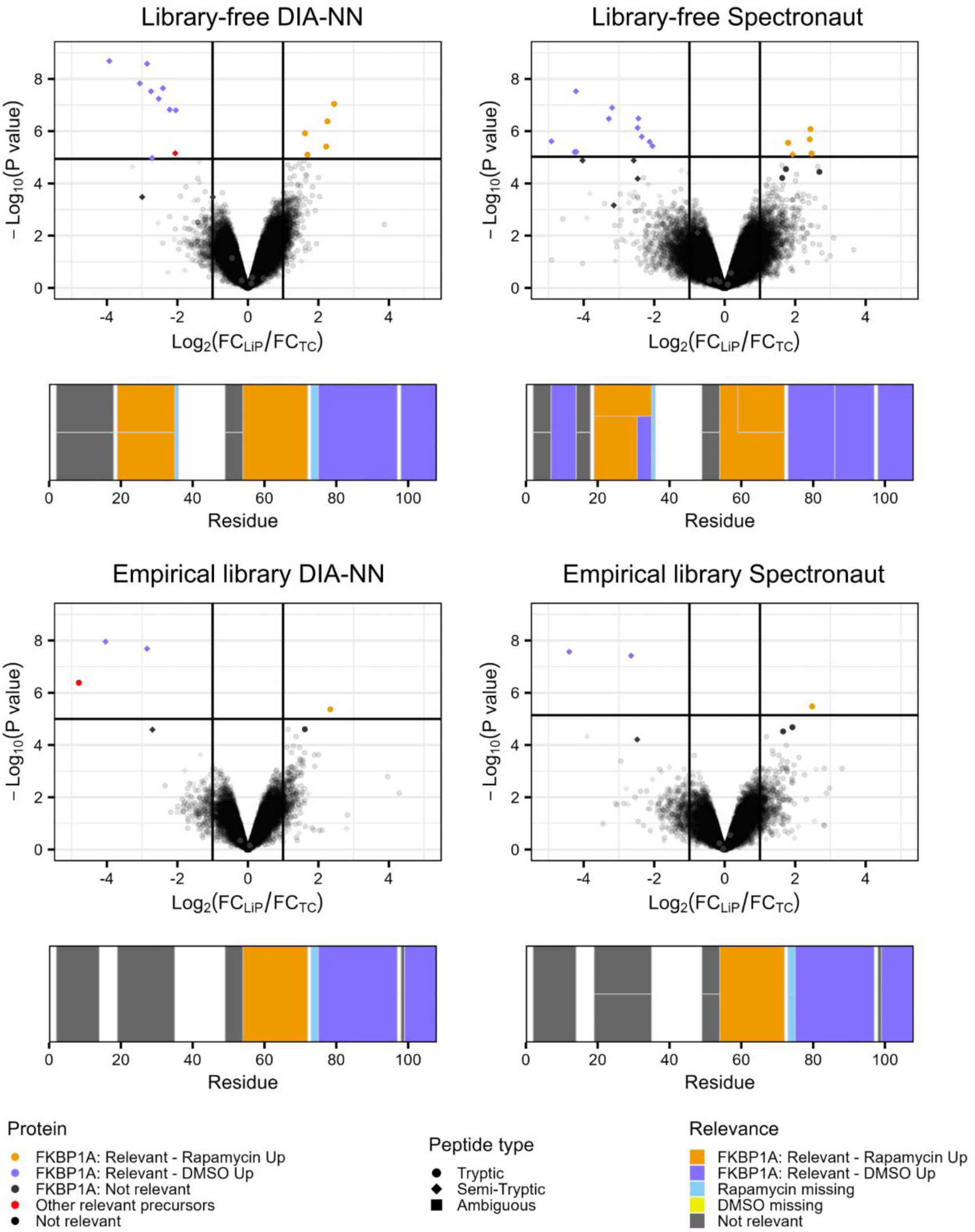
Downstream structural output of DIA LiP-MS workflows analysed with DIA-LiPA. Volcano plots show the results of the precursor-level relative differential abundance analysis between rapamycin-treated versus solvent control LiP samples for each DIA workflow. Precursors are shaped by peptide type: circles (tryptic), diamonds (semi-tryptic), squares (ambiguous/non-proteotypic peptides). Relevance thresholds: |log2(FC_LiP_/FC_TC_)| ≥ 1 and adjusted p-value ≤ 0.05. FKBP1A precursors are coloured by directionality: orange (relevant and upregulated in rapamycin-treated LiP upon correction for TC), dark blue (relevant and upregulated in solvent control LiP upon correction for TC), grey (not relevant). Other relevant hits appear in red; non-relevant precursors in black. Below each volcano plot, a structural barcode summarizes FKBP1A results for that workflow. Bars indicate precursor positions along the protein sequence, coloured by relevance: orange (relevant and upregulated in rapamycin-treated LiP upon correction for TC), dark blue (relevant and upregulated in solvent control LiP upon correction for TC), grey (not relevant), light blue (missing in rapamycin-treated), yellow (missing in solvent control), white (not detected). FC; fold change.

**Figure 4:**
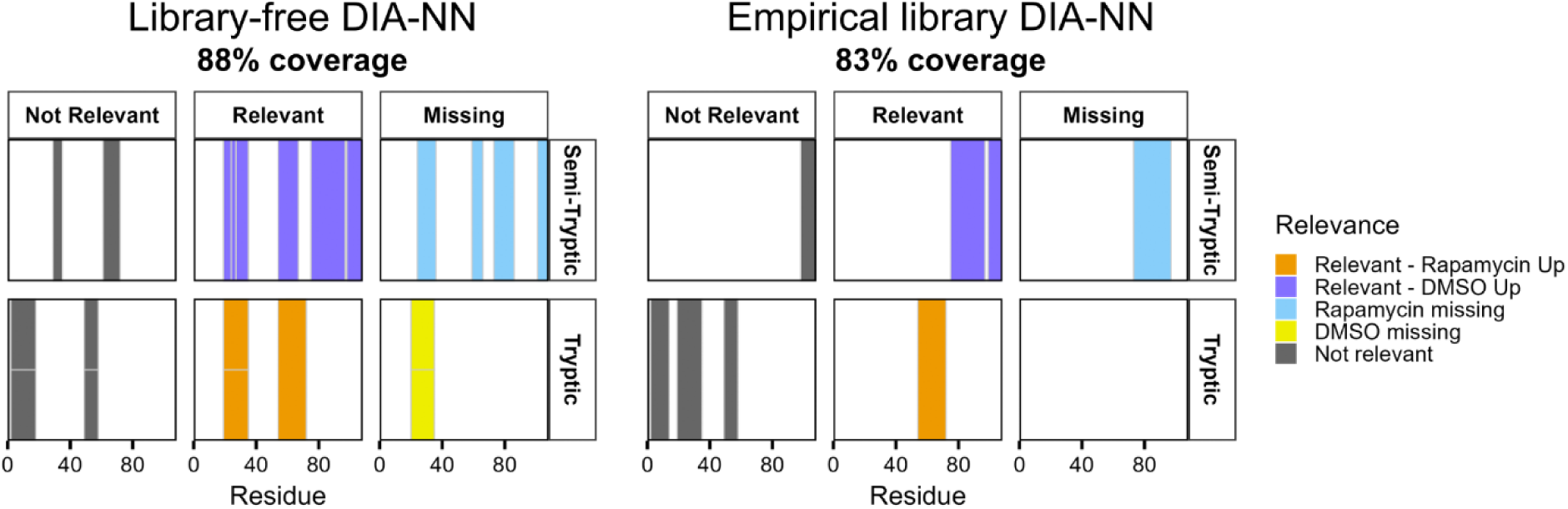
Extended structural barcodes for FKBP1A in library-free and empirical DIA-NN workflows. Each panel represents the FKBP1A sequence segmented by precursor trypticity (rows: semi-tryptic, tryptic) and detection status (columns: not relevant, relevant, missing). Relevance thresholds: |log_2_(FC_LiP_/FC_TC_)| ≥ 1 and adjusted p-value ≤ 0.05. Bars indicate precursor positions along the protein sequence, coloured by relevance: orange (relevant and upregulated in rapamycin-treated LiP upon correction for TC), dark blue (relevant and upregulated in solvent control LiP upon correction for TC), grey (not relevant), light blue (missing in rapamycin-treated), yellow (missing in solvent control), white (not detected). Bars are stacked when there are multiple precursors corresponding to the same peptide sequence. Coverage percentages for each workflow are shown above the barcodes.

### Validation of DIA-LiPA on yeast dataset

To demonstrate the generalizability and utility of DIA-LiPA, we reanalysed a publicly available yeast heat shock dataset ^31^ using two library-free workflows (DIA-NN and Spectronaut), followed by downstream analysis via DIA-LiPA. Because this dataset contains inherent sample pairing between LiP and TC, we used our paired DIA-LiPA pipeline.

Precursor identifications and overlaps between workflows demonstrate high consistency and comprehensive coverage, with both tryptic and semi-tryptic peptides contributing substantially (**Figure 5A**). Semi-tryptics represent roughly 38% of all identifications, indicative of a dataset rich in structural information.

The structural barcode of DIA-NN for Hsp104 (**Figure 5B**), a key disaggregase highlighted in the original study, reveals domain-level regulation patterns. While both search engines detect substantially more precursors and thereby provide improved coverage relative to the original study, only the DIA-NN barcode reached statistical significance, while Spectronaut did not. This difference arises because Spectronaut detects and retains a larger number of precursors, leading to more statistical tests and therefore a more stringent multiple-testing correction, which yields higher adjusted p-values (**Supplementary Figure S6A**). Together with our previous entrapment-based FDR assessment, these results indicate that Spectronaut may require more stringent filtering. Because of this, only the DIA-NN barcode provides a statistically supported structural read-out that aligns with the observations of Cappelletti *et al.*, who mapped altered peptides to ATP-binding sites, substrate channels and solvent-exposed regions and suggested that multiple structural states coexist in the sample. While their fractionation approach (soluble versus insoluble) was designed to disentangle these states, our DIA-LiPA analysis of total lysates alone recapitulates these insights and adds clarity on domain-specific changes. For example, the expanded barcode (**Supplementary Figure S7**) demonstrates that the middle region (MD) contains mostly semi-tryptics upregulated after heat shock upon correction for TC, consistent with a region that is simultaneously flexible, supporting its role in modulating substrate threading and ATPase activity during heat shock ^48,49^. In addition to the middle domain, multiple semi-tryptic peptides in both the N- and C-terminal domains showed upregulation after heat shock upon correction for TC, indicating enhanced accessibility and domain-level flexibility. This pattern is consistent with the conformational activation cycle of Hsp104, where the N- and C-terminal regions undergo dynamic rearrangements to support substrate engagement and hexamer restructuring ^50,51^, reinforcing the notion that multiple structural states can coexist.

Volcano plots are a common tool for the visualisation of the results from differential abundance analyses. In LiP-MS they can provide a global view of structural dynamics by regarding tryptic and semi-tryptic peptides separately (**Figure 5C**). Tryptic precursors exhibit a pronounced shift toward negative relative fold changes, upon adjustment for protein-level fold changes in the trypsin control samples. The trend of decreased tryptic peptide abundance upon correction for TC likely reflects increased proteolytic accessibility due to unfolding, making more sites surface accessible, leading to more aspecific cleavage events and consequently lowering tryptic signal. In an ideal detection scenario, semi-tryptic counterparts of these peptides would be consistently observed and exhibit a complementary shift toward positive fold changes upon correction for TC. This is precisely the pattern expected when previously buried regions become exposed. Indeed, for the semi-tryptic precursors that are detected in both conditions, we observe this expected positive shift, supporting the interpretation of heat-induced structural opening. However, volcano plots only include precursors for which a statistical comparison is possible, i.e. those with detectable values in both conditions. Semi-tryptic peptides arising from regions that become exposed only upon heat shock are absent in the control condition and therefore cannot be assigned a p-value and are thus absent in the plot. This limitation is especially relevant for semi-tryptic peptides originating from regions such as buried hydrophobic cores that become surface-accessible only after heat-induced unfolding. This highlights the added value of integrating missingness patterns to detect structural changes beyond the protein surface. Our workflow further supports both absolute (abundance-based) and relative-abundance (TC-adjusted) statistical analyses, which can reveal distinct aspects of structural change (**Supplementary Figure S6B**). In the yeast dataset, applying TC-adjustment substantially changed the differential abundance profile, demonstrating that relative-abundance analysis can robustly correct for protein-level fold changes, revealing site- or region-specific effects.

Together, these findings confirm that DIA-LiPA enhances interpretability of existing datasets and uncovers additional structural insights, supporting simple and complex experimental designs with both paired or unpaired LiP and TC samples.

**Figure 5:**
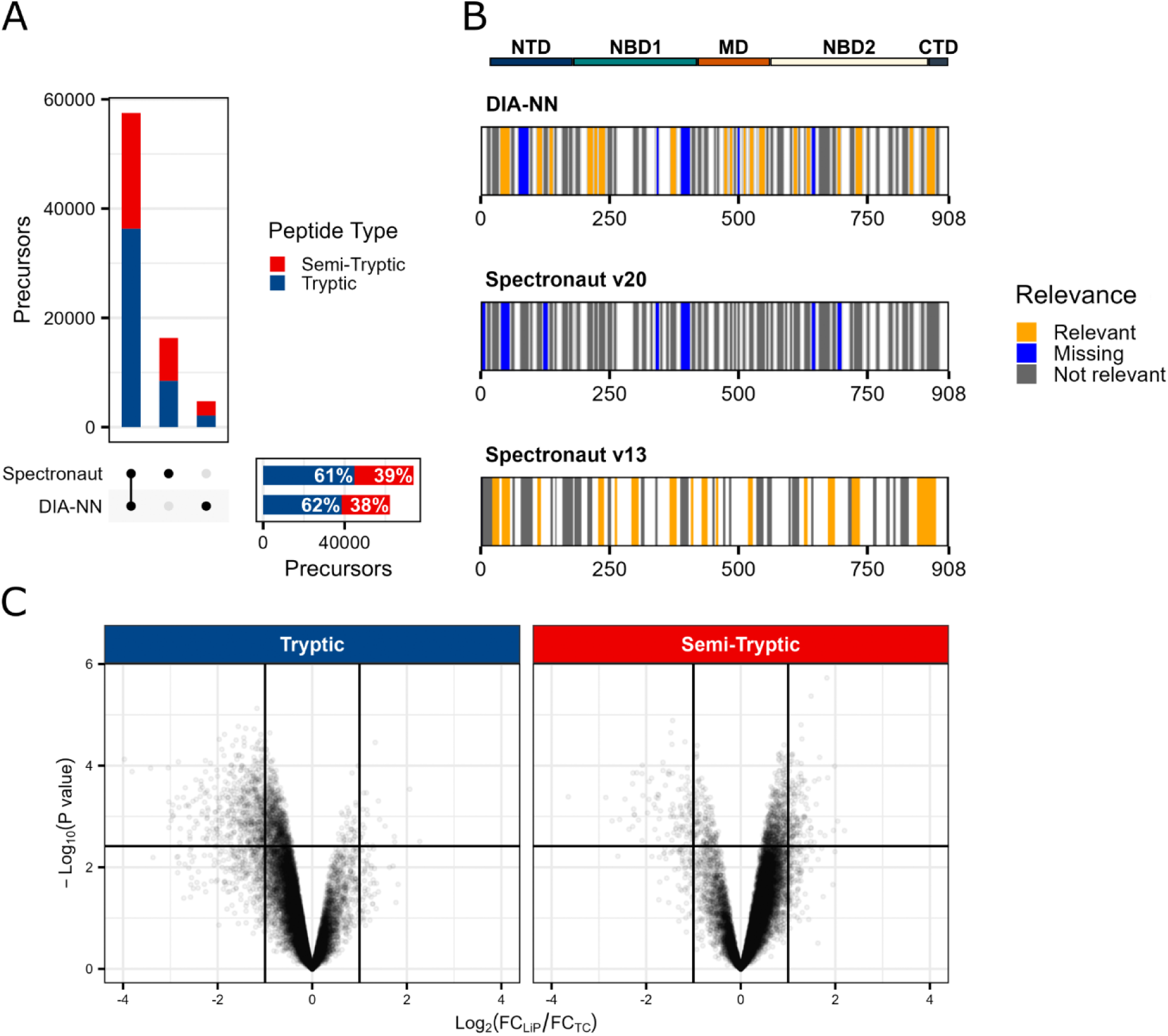
Validation of DIA-LiPA on the yeast heat shock dataset. **A)** UpSet plot showing the absolute number of precursors in LiP samples identified by library-free DIA-NN and Spectronaut workflows and their overlaps. Percentages of tryptic and semi-tryptic precursors are indicated for each workflow. Only post-filtered data are shown (q-value < 0.01, each precursor was identified at least twice in at least one condition). **B)** Structural barcodes of Hsp104, a highlighted target in the original study, for both workflows alongside the original barcode from the study. Relevance thresholds: |log_2_(FC_LiP_/FC_TC_)| ≥ 1 and adjusted p-value ≤ 0.05. Each vertical bar represents a potential LiP precursor, coloured by status: orange (relevant), blue (missing), grey (not relevant), white (not detected by MS). Bars are stacked when there are multiple precursors corresponding to the same peptide sequence. Protein domains (N-terminal domain; NTD, nucleotide-binding domain 1; NBD1, middle domain; MD, nucleotide-binding domain 2; NBD2, C-terminal domain; CTD) are indicated on top. **C)** Volcano plots of library-free DIA-NN results separated by precursor trypticity. Left: tryptic precursors; right: semi-tryptic precursors. Each point represents a precursor. Relevance thresholds are indicated by vertical and horizontal lines: |log_2_(FC_LiP_/FC_TC_)| ≥ 1 and adjusted p-value ≤ 0.05. Only proteins that contain both tryptic and semi-tryptic precursors in the differential abundance output are shown.

### DIA-LiPA can also be applied to PELSA data analysis

In addition to validating DIA-LiPA on LiP-MS datasets, we examined whether the pipeline can accommodate related limited proteolysis approaches such as PELSA ^10^. Using the publicly available rapamycin PELSA dataset (PXD034606), we processed the data with a minimally adapted version of DIA-LiPA. Because the experimental design employed in this study does not include trypsin controls, the analysis reduces to a standard differential abundance workflow. Note that PELSA relies solely on long, frequently miscleaved, fully tryptic peptides and therefore does not provide information on semi-tryptic LiP fragments, thus limiting its structural resolution. Regardless, our results reproduced the protein-level shifts reported in the original study (**Supplementary Figure S8**), while the absence of semi-tryptic peptides precluded the detailed structural interpretation achievable with LiP-MS. This demonstrates that the DIA-LiPA workflow is not strictly limited to classical LiP-MS data.

## Conclusions

Our findings illustrate how recent advances in machine learning have transformed DIA workflows, making library-free approaches not only sufficient but potentially superior to traditional DDA-based libraries. This shift reduces experimental overhead, instrument time and sample requirements, critical advantages for applications with limited material such as clinical biopsies.

In this context, we introduce DIA-LiPA, a pipeline tailored for classical LiP-MS and suitable for PELSA that integrates semi-tryptic peptides and uniquely accounts for missingness to enable a more nuanced structural interpretation, revealing subtle conformational changes with high confidence. These capabilities extend beyond peptide quantification, offering a framework for mechanistic insights into protein dynamics.

While benchmarking DIA-NN’s new semi-specific digestion option confirms its robustness, the broader impact of this work lies in demonstrating how DIA-LiPA leverages these advances to deliver reproducible, high-resolution structural information. Validation across two datasets, namely the rapamycin-treated human cell lysate and yeast heat shock, shows that DIA-LiPA not only reproduces known structural signatures but also uncovers additional layers of regulation. Importantly, DIA-LiPA can explicitly adjust for overall protein level changes in tryptic controls (TC), which can either be paired or unpaired with the LiP samples; and correctly propagates the uncertainty in the protein-level TC quantities in the downstream relative differential abundance report.

Together, these findings position DIA-LiPA as a valuable addition to the LiP-MS toolbox and a step toward more accessible, high-confidence structural proteomics. Furthermore, we also provide an alternative search strategy including a semi-tryptic FASTA file which may be adapted based on the provided scripts to serve applications outside of LiP-MS.

## Author Contributions

All authors contributed to the final version of the manuscript. ‡CVL and EA contributed equally to this work.

## Notes

The authors declare no competing financial interest.

## Acknowledgement

We acknowledge the VIB Proteomics Core for their support in performing the mass spectrometry measurements. We thank Ludovic Gillet and Paola Picotti (ETH Zurich) for providing initial expertise and guidance on the LiP-MS protocol. We also thank Vadim Demichev for his valuable feedback on FDR validation, which helped refine the interpretation of entrapment results in this study. Finally, we thank the Biognosys support team for their helpful technical assistance regarding raw-file handling and the implementation of the Semi-Specific Pipeline with Full Enumeration in Spectronaut.

## Footnotes

1 https://github.com/Gevaert-Lab/dialipar

2 https://github.com/vdemichev/DiaNN/releases/tag/2.0

3 https://github.com/Gevaert-Lab/dialipar

