## Supporting Information for "Advancing DIA-based Limited Proteolysis Workflows: introducing DIA-LiPA"

**Contents:**

Supplementary methods section.

Figure S1: Precursor q-value and intensity distributions by workflow.

Figure S2: Coverage and CV distributions for non-shared precursors across DIA LiP-MS workflows.

Figure S3: Benchmarking of semi-tryptic analysis strategies in DIA-NN.

Figure S4: Representative FKBP1A spectra from DIA-NN and Spectronaut.

Figure S5: Extended structural barcodes for FKBP1A in library-free and empirical Spectronaut workflows.

Figure S6: Effect of search engine and relative abundance modeling in the yeast dataset.

Figure S7: Extended structural barcode for Hsp104 in library-free DIA-NN.

Figure S8: Re-analysis of PELSA data.

Table S1: Impact of HTRMS conversion and Semi-Specific Pipeline settings on semi-tryptic identifications in Spectronaut 20.3.

#### SUPPLEMENTARY METHODS SECTION

##### The semi-tryptic FASTA workaround for DIA-NN (version 2.2.0)

A semi-tryptic peptide FASTA file was generated from the human UniProt FASTA (01/2024) with the proteinase K (*Tritirachium album*) sequence appended, using a custom R script (available on MassIVE (MSV000099740): folder sequence/SemiTrypticFasta_generation). Proteins were *in silico* digested to include both fully and semi-tryptic sequences ranging from 7 to 30 amino acids in length, allowing up to one missed cleavage. All peptides generated this way were used to construct a peptide FASTA file, with headers derived from the protein, retaining the original protein header to ensure consistent annotation. Next, this peptide FASTA file was used as input for spectral library prediction in DIA-NN. The missed cleavages parameter was set to 0, preventing DIA-NN from generating additional missed cleavage variants, which would be redundant. The resulting predicted library was then used to search the DIA raw files. The following additional options were specified: *--cut*, to disable further *in silico* digestion, and *--duplicate-proteins*, to retain entries with duplicate FASTA headers.

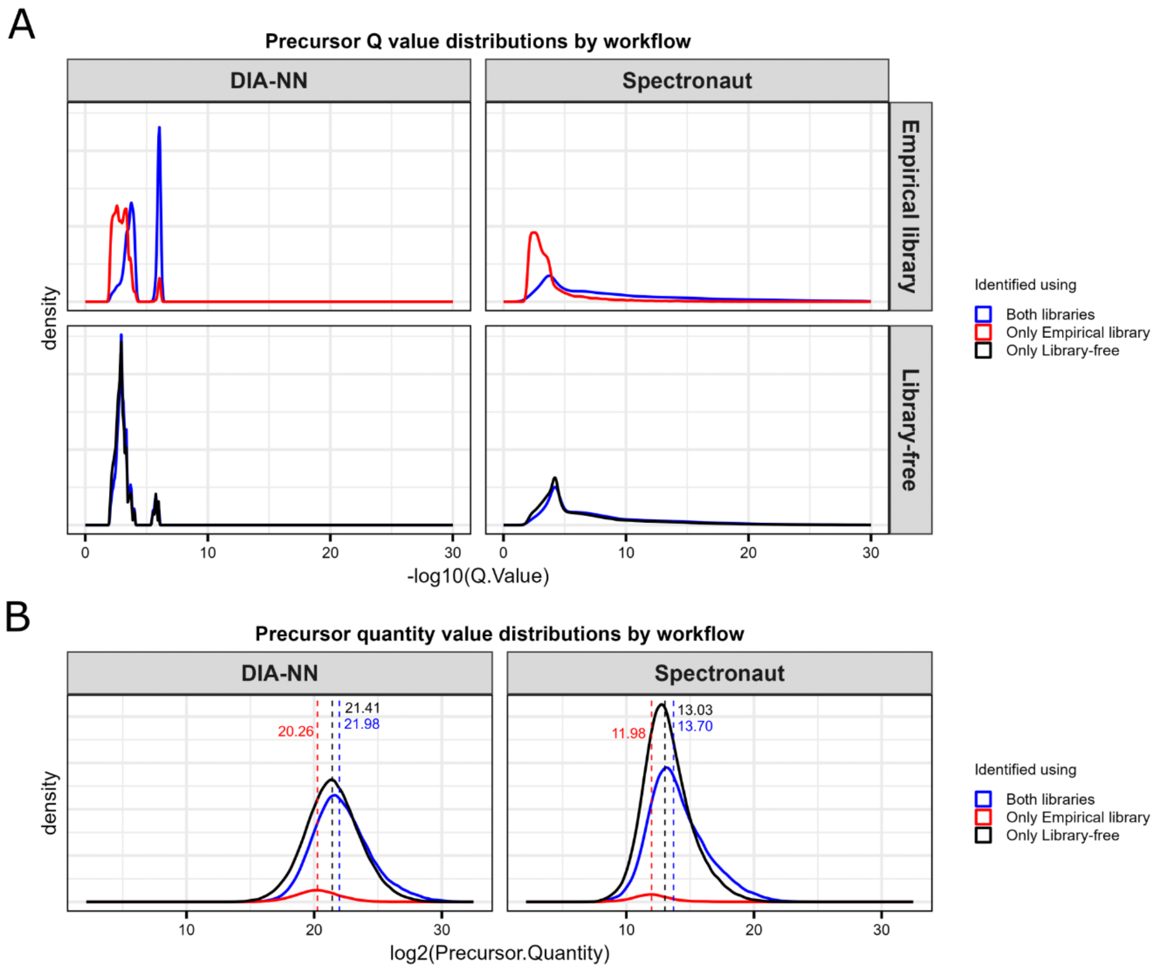


**Figure S1: Precursor q-value and intensity distributions by workflow. A)** Distribution of precursor q-values by workflow. Traces show the density of -$\log_{10}$ precursor q-values of all precursors which were retained after filtering as implemented in the DIA-LiPA workflow. **B)** The distribution of $\log_2$ precursor intensities identified by each workflow. Traces depict the $\log_2$ MS2-based intensity of precursors which were retained after filtering as part of the DIA-LiPA workflow, rescaled for the number of identifications in each category with median values indicated.

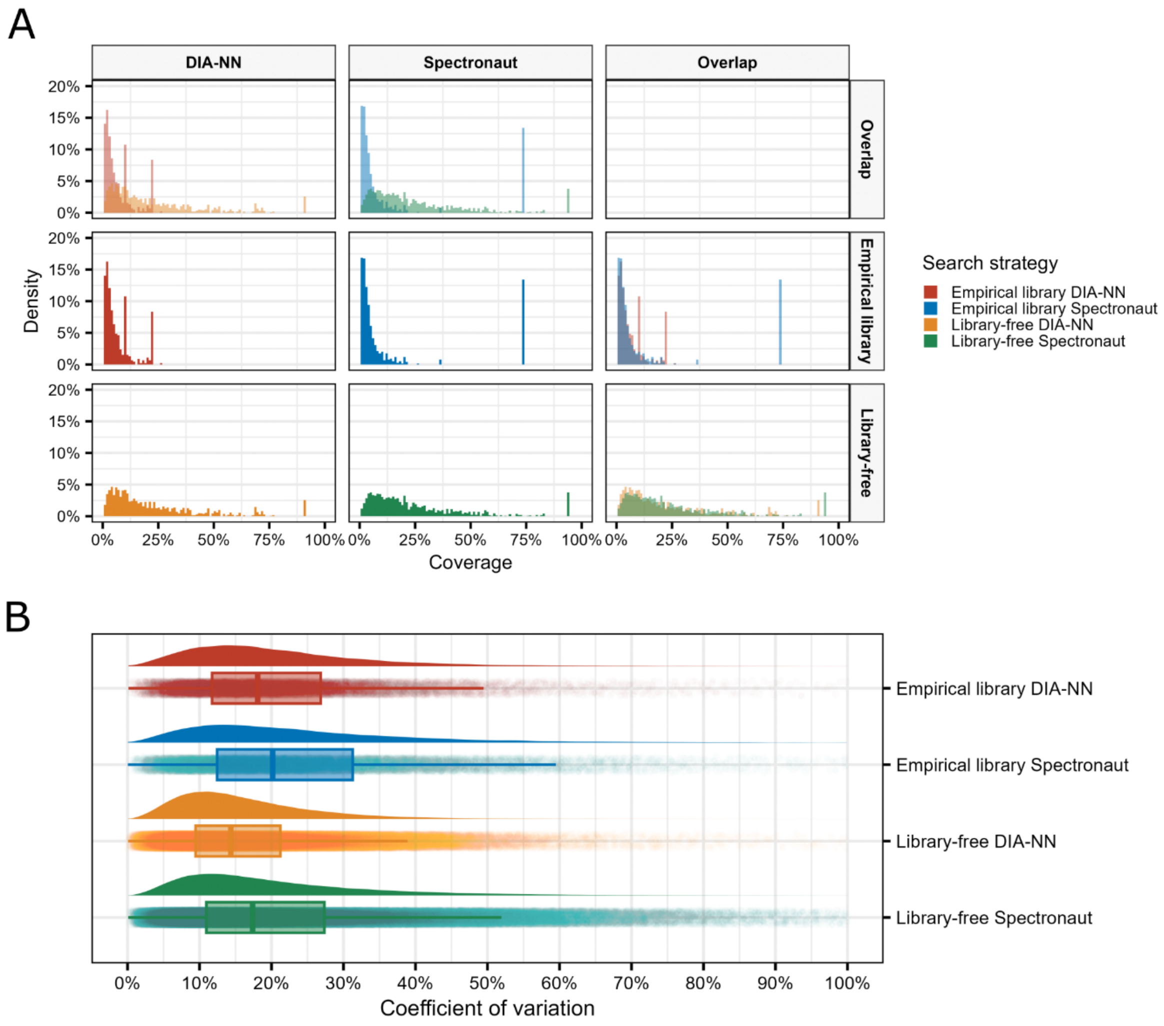


**Figure S2: Coverage and CV distributions for non-shared precursors across DIA LiP-MS workflows. A)** Protein sequence coverage distribution for non-shared proteins accessions in LiP samples across workflows and their overlaps. **B)** Raincloud plot showing the distribution of normalised precursor-level CVs for non-shared precursors for each workflow. The density curve represents the CV distribution, while the boxplot below indicates the median (line), interquartile range (box) and whiskers spreading to 1.5x the interquartile range. Individual points represent precursor-level CVs. CVs were calculated across all LiP samples per condition (rapamycin or vehicle control) for precursors quantified in at least three samples. CV; coefficient of variation.

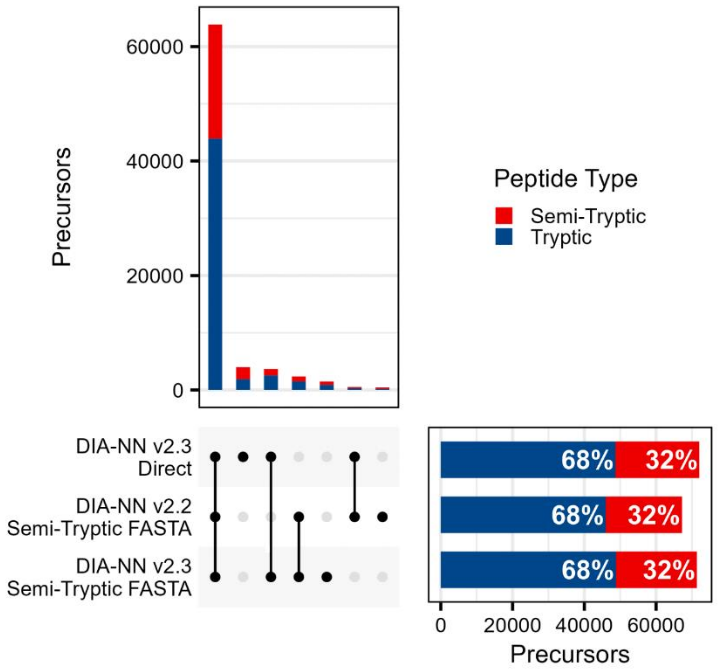


**Figure S3: Benchmarking of semi-tryptic analysis strategies in DIA-NN.** Comparison of identification rates across three DIA-NN workflows in LiP samples: semi-tryptic FASTA-based workaround in v2.2, semi-tryptic FASTA-based workaround in v2.3, and semi-specific digestion in v2.3. UpSet plot showing the absolute number of precursors identified by each workflow and their overlaps. Percentages of tryptic and semi-tryptic precursors are indicated for each workflow. Only post-filtered data are shown (q-value < 0.01, each precursor was identified at least twice in at least one condition).

**Figure S4: Representative spectra for FKBP1A precursors identified in the rapamycin LiP-MS experiment.** Representative tryptic and semi-tryptic FKBP1A precursors detected by library-free DIA-NN (v2.3.0) are shown with chromatographic elution profiles and annotated fragment-ion spectra visualized in Skyline (pages 7, 9, 11, 13, 15, 17). On alternating pages, the corresponding precursors identified by Spectronaut (v20.3) are shown using its native spectrum viewer (pages 8, 10, 12, 14, 16, 18). Generally, only the top six fragment ions are displayed, as annotated by Skyline or Spectronaut.

For each precursor, the upper panel displays the elution profiles across all samples, and the lower panel shows the annotated fragment-ion intensities alongside the predicted spectrum.

### AKLTISPDYAYGAT2

Semi-tryptic; present in DMSO LiP only

DIA-NN v2.3, visualization in Skyline

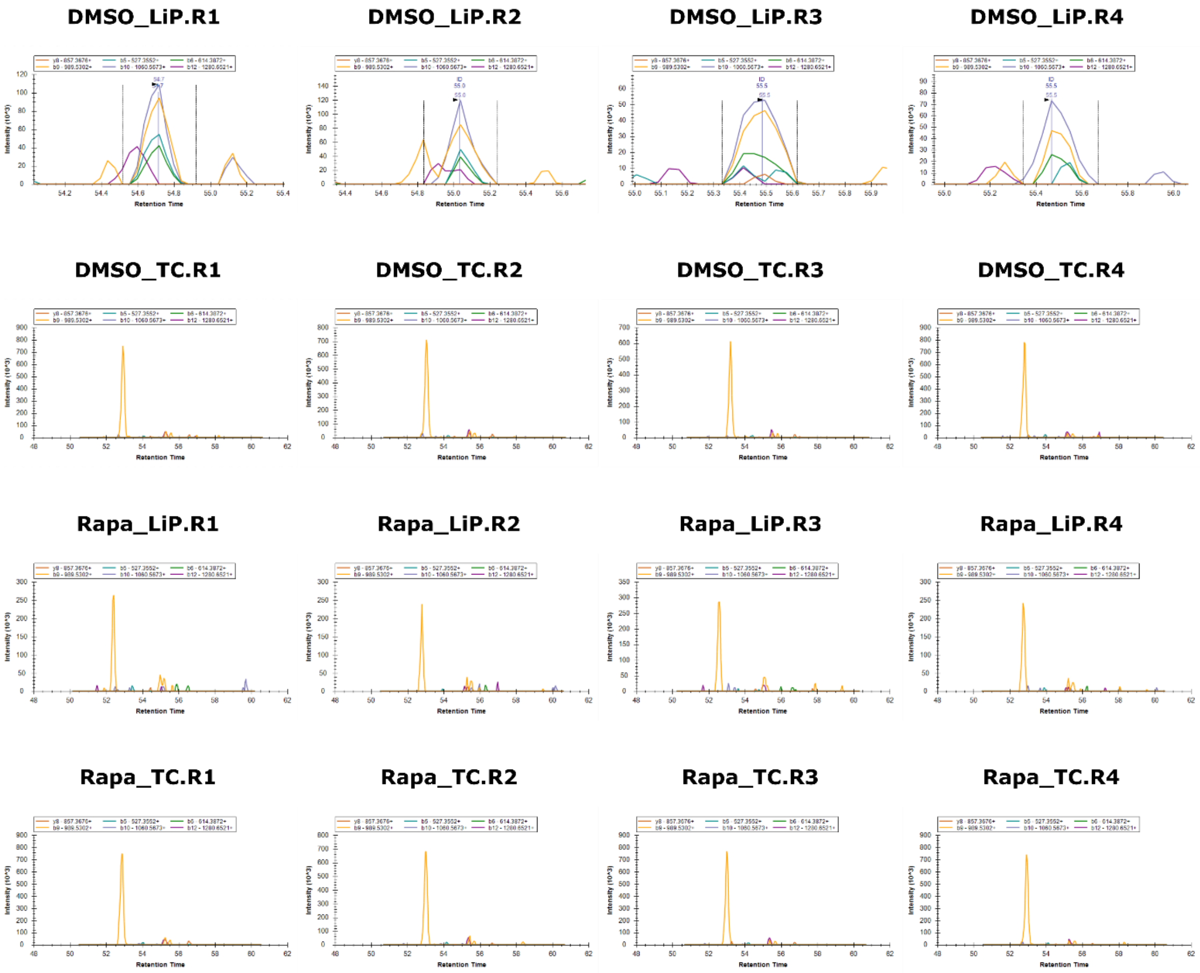


#### Annotated versus predicted spectrum

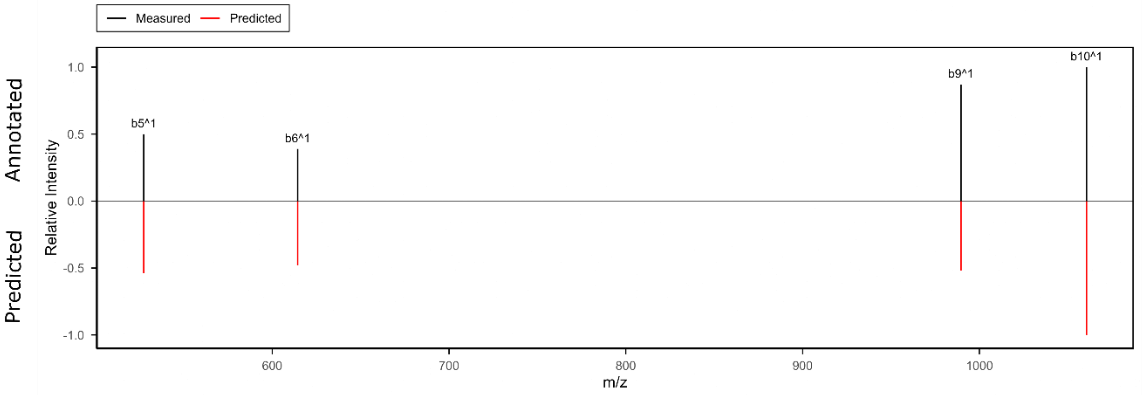

### AKLTISPDYAYGAT2

Semi-tryptic; present in DMSO LiP only

Spectronaut v20.3

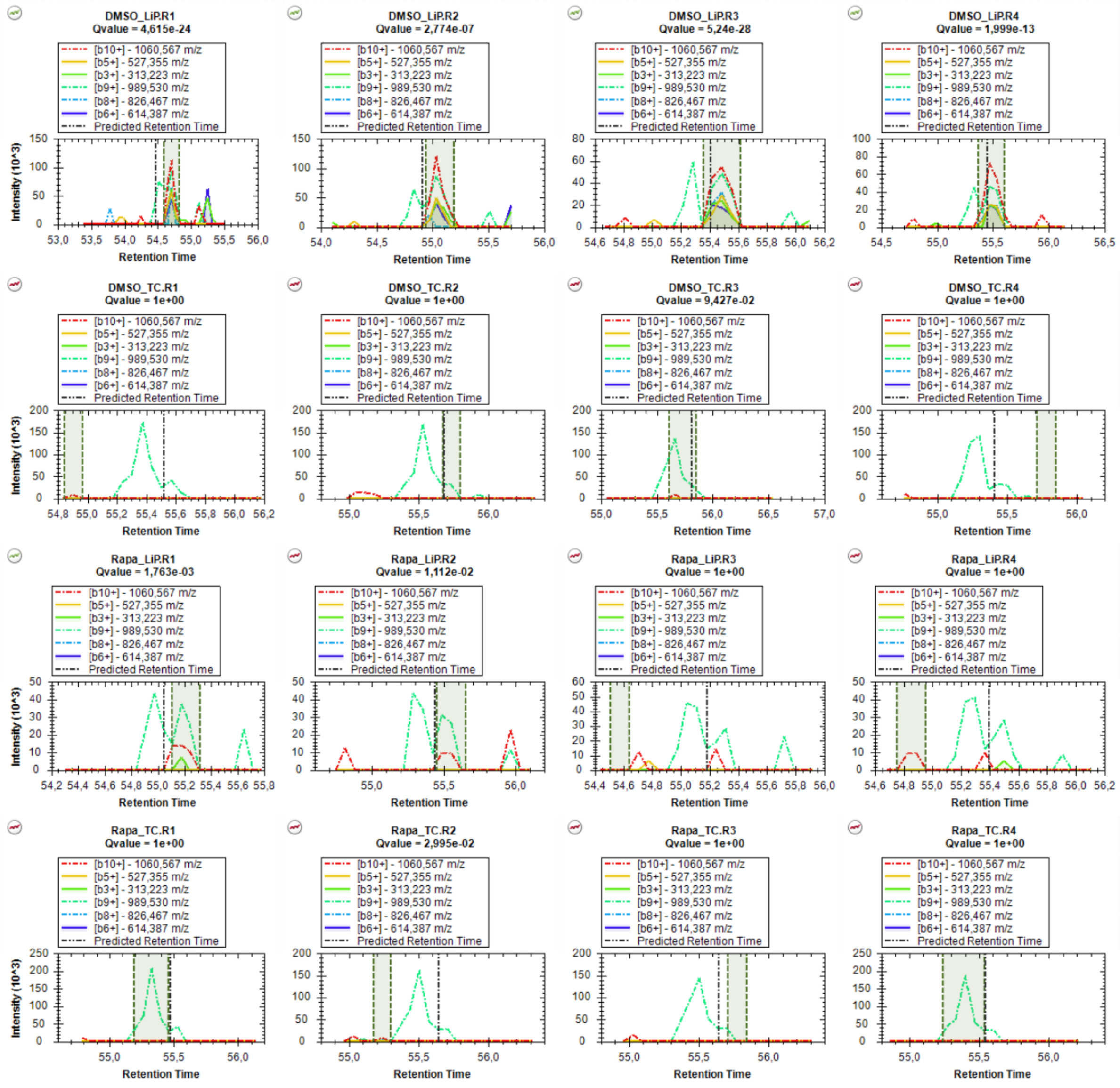


#### Annotated versus predicted spectrum

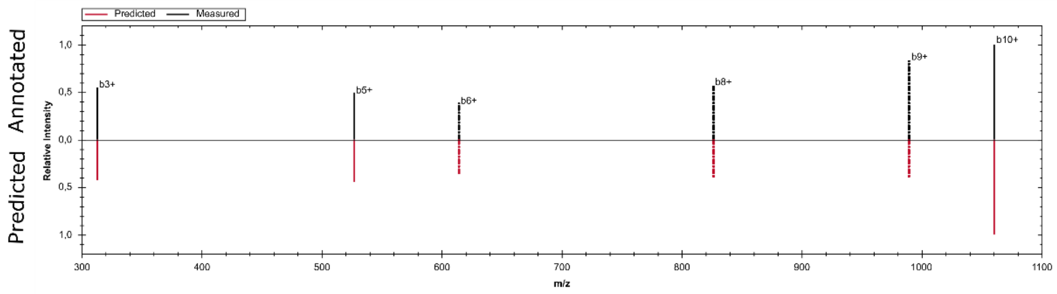

### GWEEGVAQMSVGQR2

Tryptic; present in all conditions

DIA-NN v2.3, visualization in Skyline

**DMSO_LiP.R1**

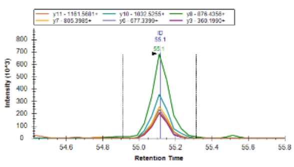

**DMSO_LiP.R2**

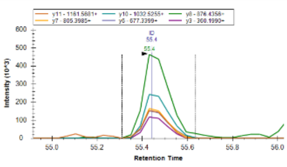

**DMSO_LiP.R3**

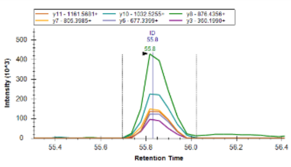

**DMSO_LiP.R4**

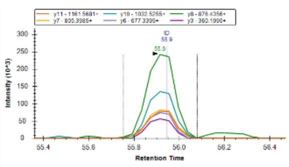

**DMSO_TC.R1**

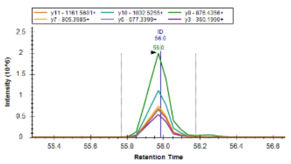

**DMSO_TC.R2**

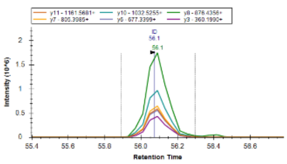

**DMSO_TC.R3**

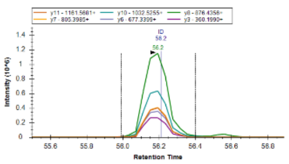

**DMSO_TC.R4**

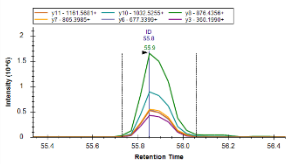

**Rapa_LiP.R1**

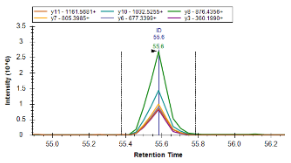

**Rapa_LiP.R2**

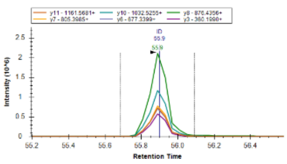

**Rapa_LiP.R3**

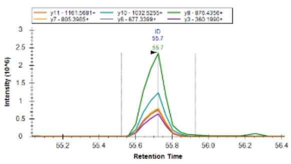

**Rapa_LiP.R4**

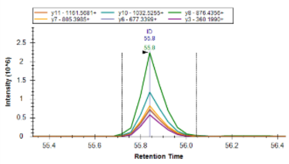

**Rapa_TC.R1**

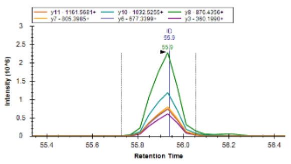

**Rapa_TC.R2**

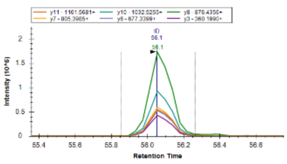

**Rapa_TC.R3**

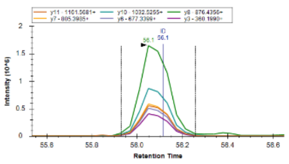

**Rapa_TC.R4**

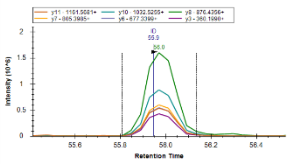

**Annotated versus predicted spectrum**

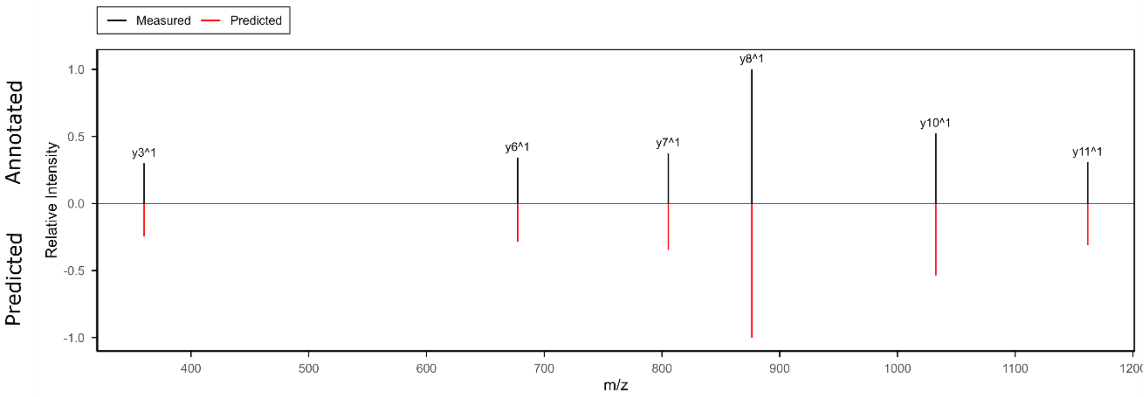

### GWEEGVAQMSVGQR2

Tryptic; present in all conditions

Spectronaut v20.3

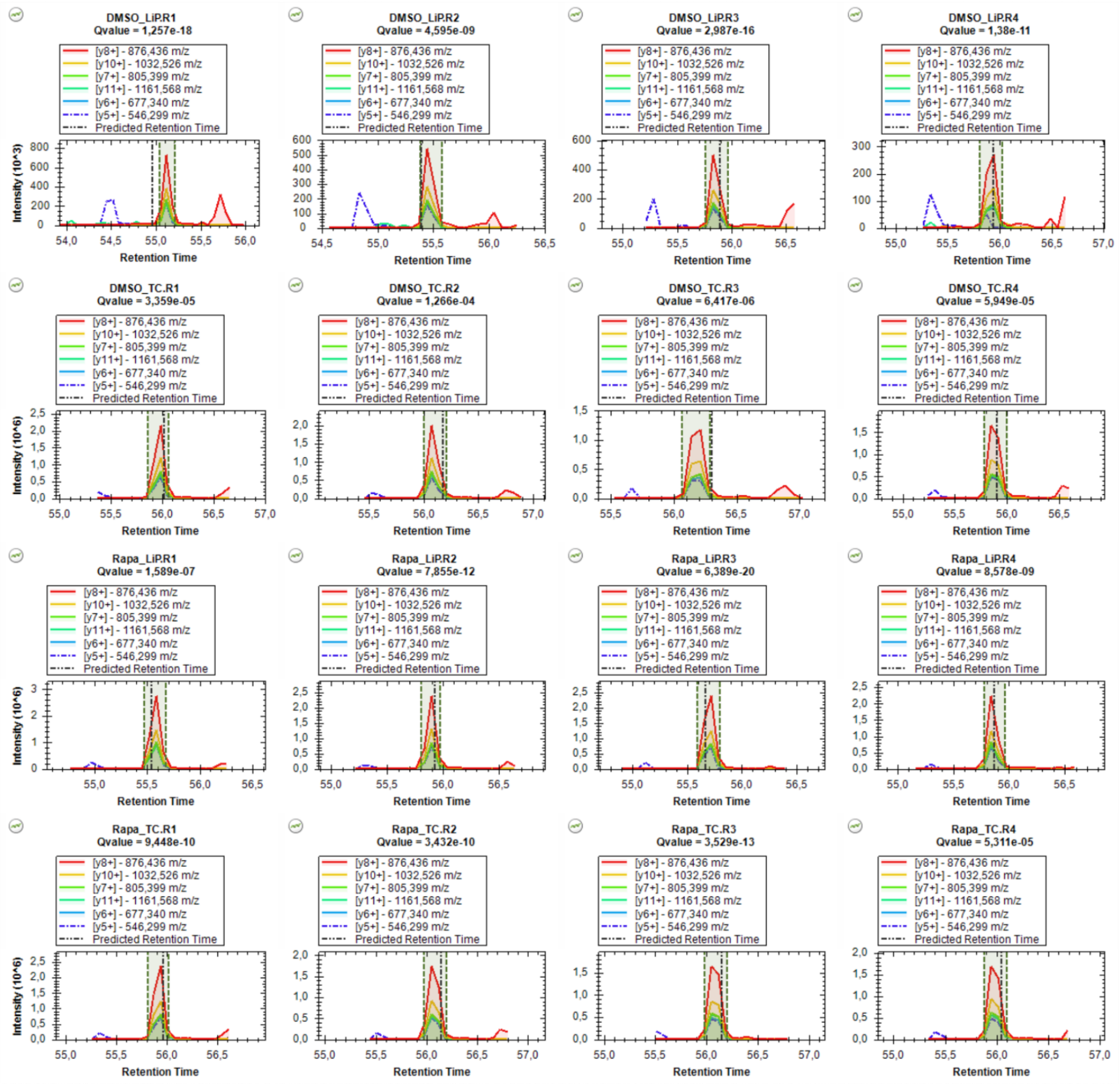


#### Annotated versus predicted spectrum

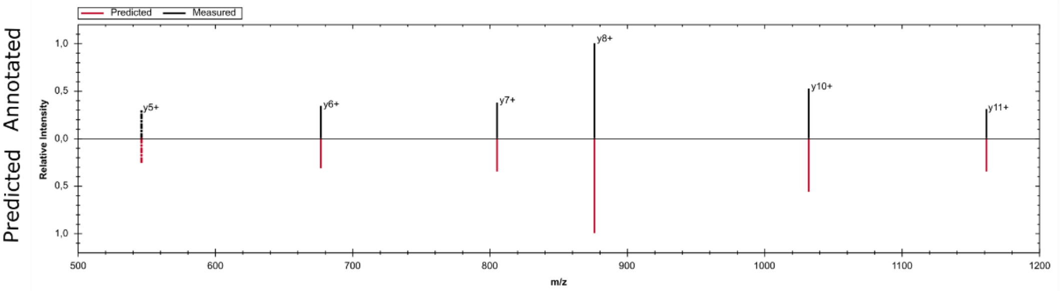

### LVFDVELLK2

Semi-tryptic; present in LiP conditions

DIA-NN v2.3, visualization in Skyline

**DMSO_LiP.R1**

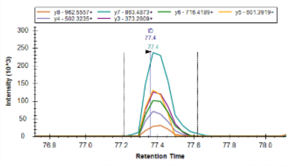

**DMSO_LiP.R2**

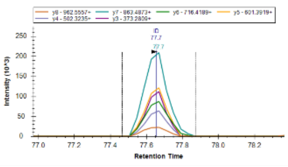

**DMSO_LiP.R3**

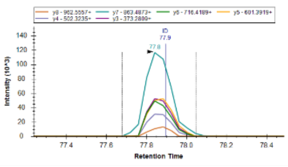

**DMSO_LiP.R4**

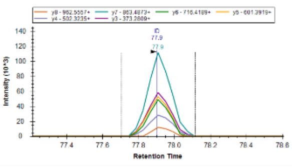

**DMSO_TC.R1**

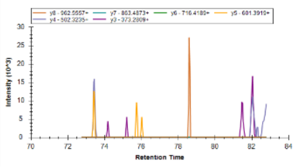

**DMSO_TC.R2**

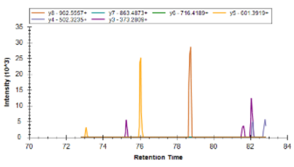

**DMSO_TC.R3**

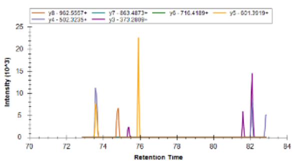

**DMSO_TC.R4**

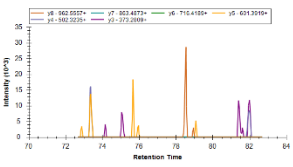

**Rapa_LiP.R1**

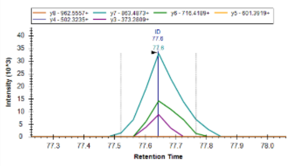

**Rapa_LiP.R2**

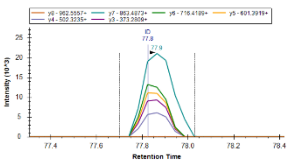

**Rapa_LiP.R3**

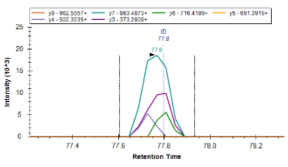

**Rapa_LiP.R4**

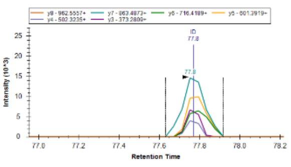

**Rapa_TC.R1**

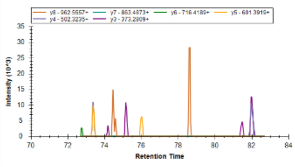

**Rapa_TC.R2**

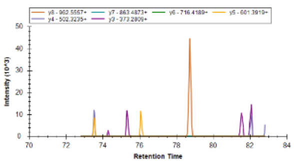

**Rapa_TC.R3**

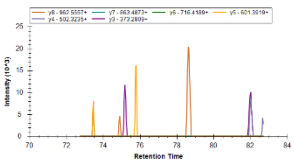

**Rapa_TC.R4**

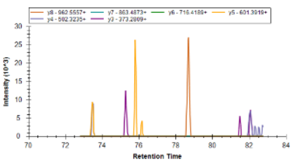

**Annotated versus predicted spectrum**

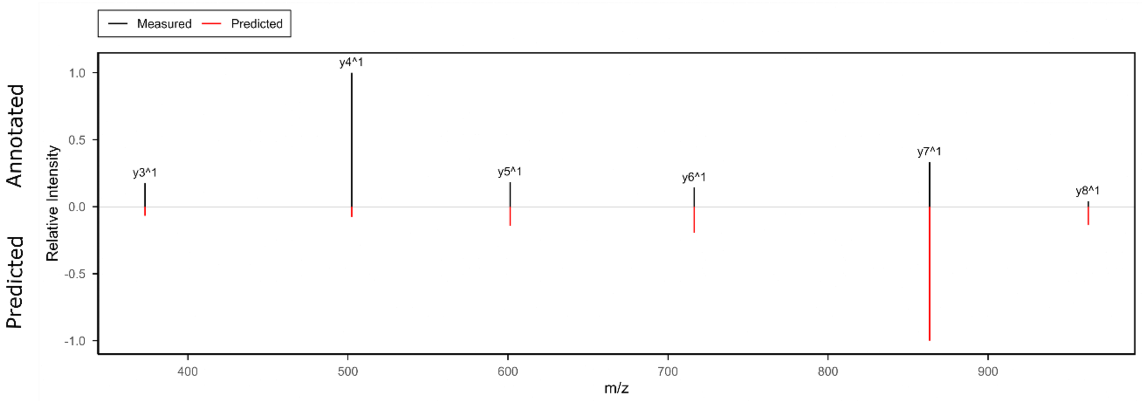

### LVFDVELLK2

Semi-tryptic; present in LiP conditions

Spectronaut v20.3

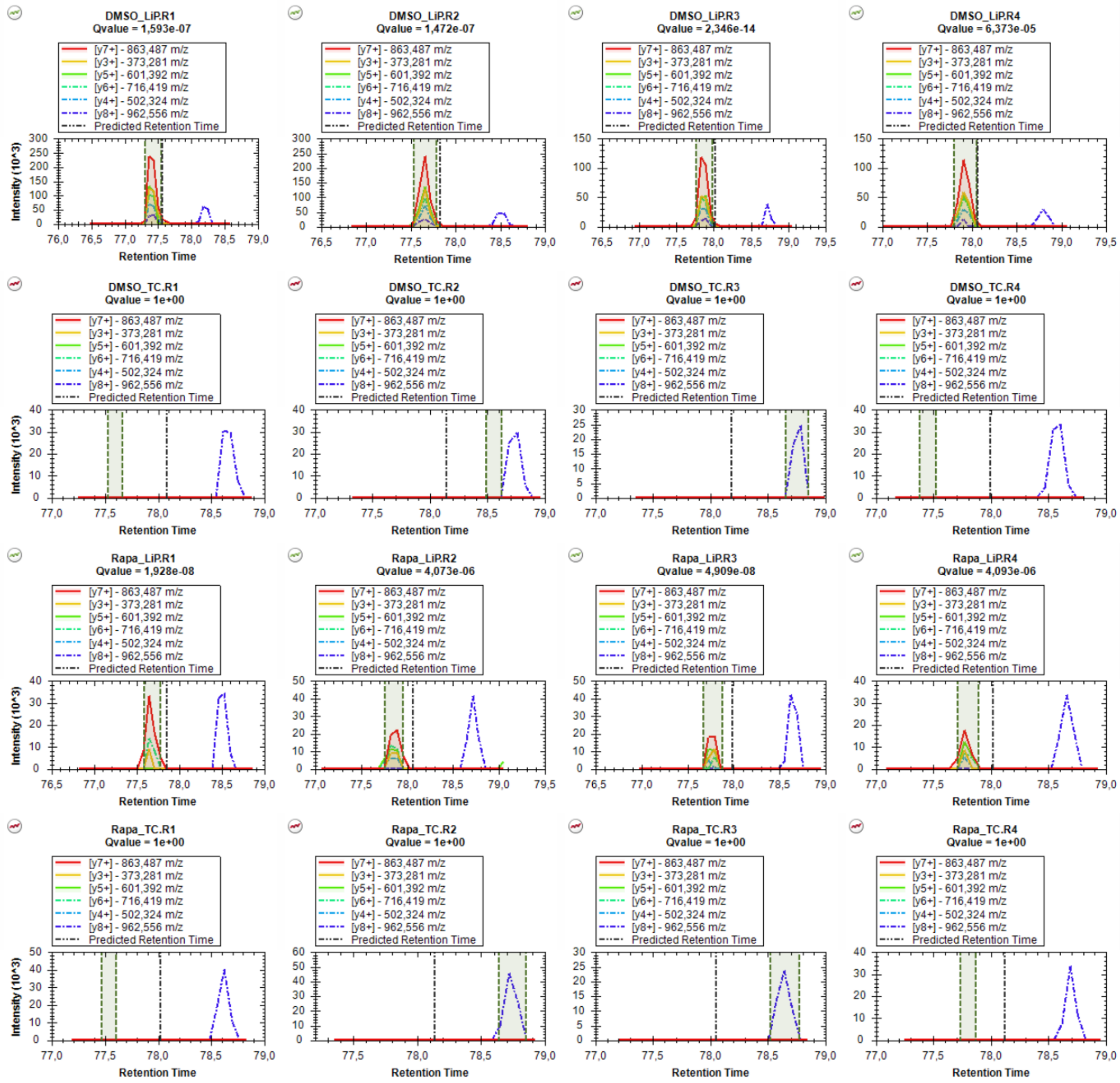


#### Annotated versus predicted spectrum

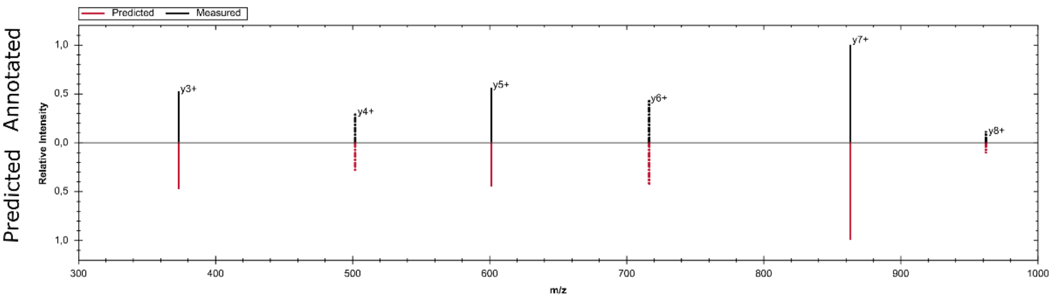

### YTGMLEDGK2

Semi-tryptic; present in LiP conditions

DIA-NN v2.3, visualization in Skyline

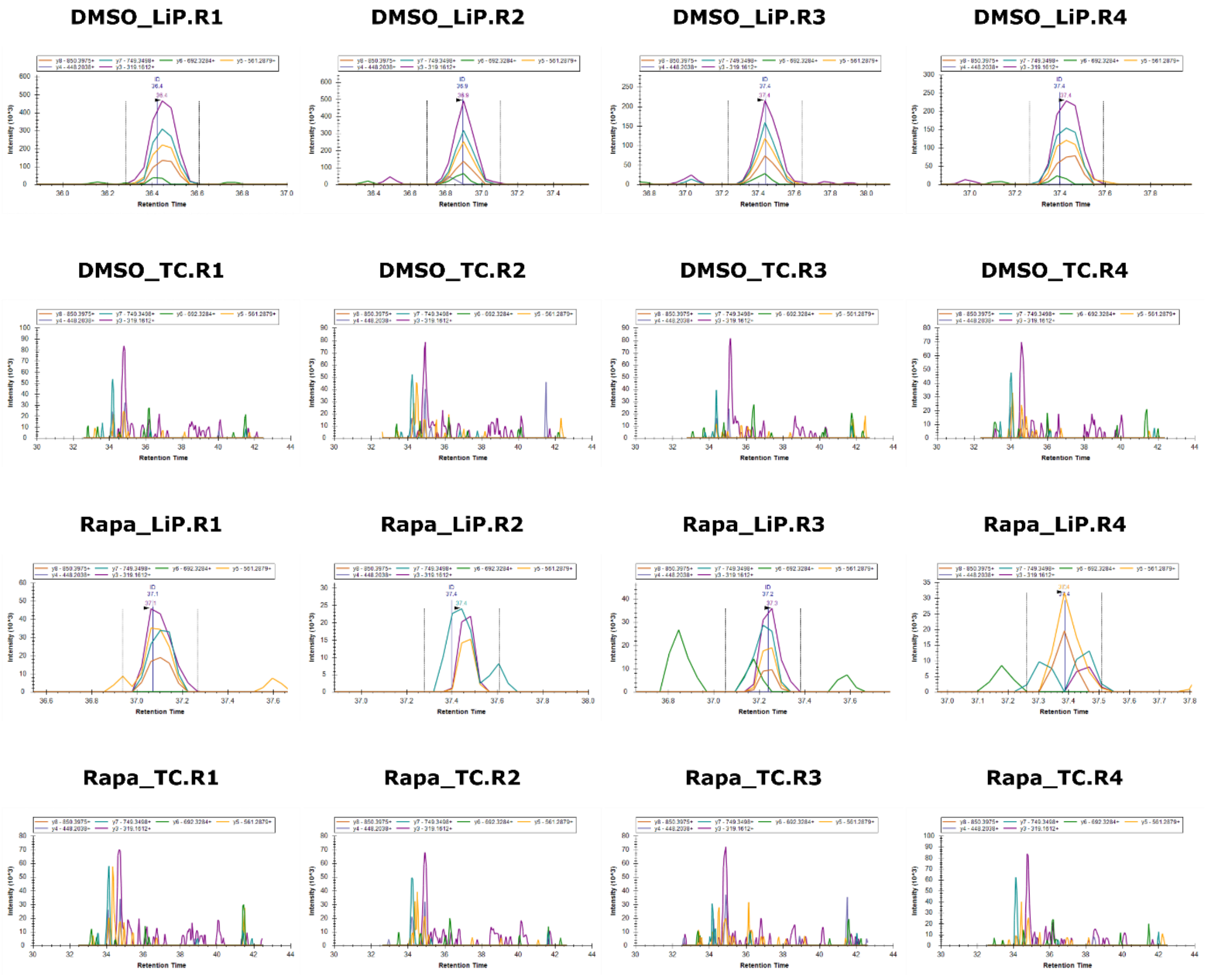


#### Annotated versus predicted spectrum

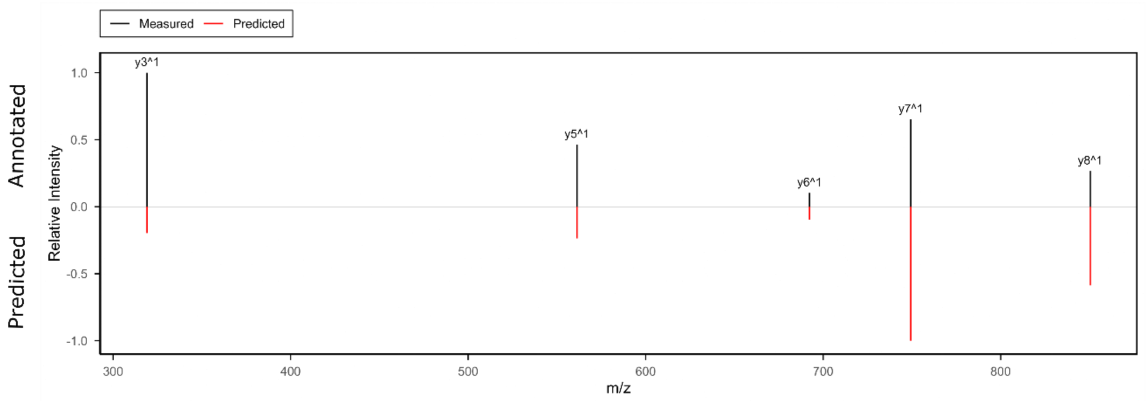

### YTGMLEDGK2

Semi-tryptic; present in LiP conditions

Spectronaut v20.3

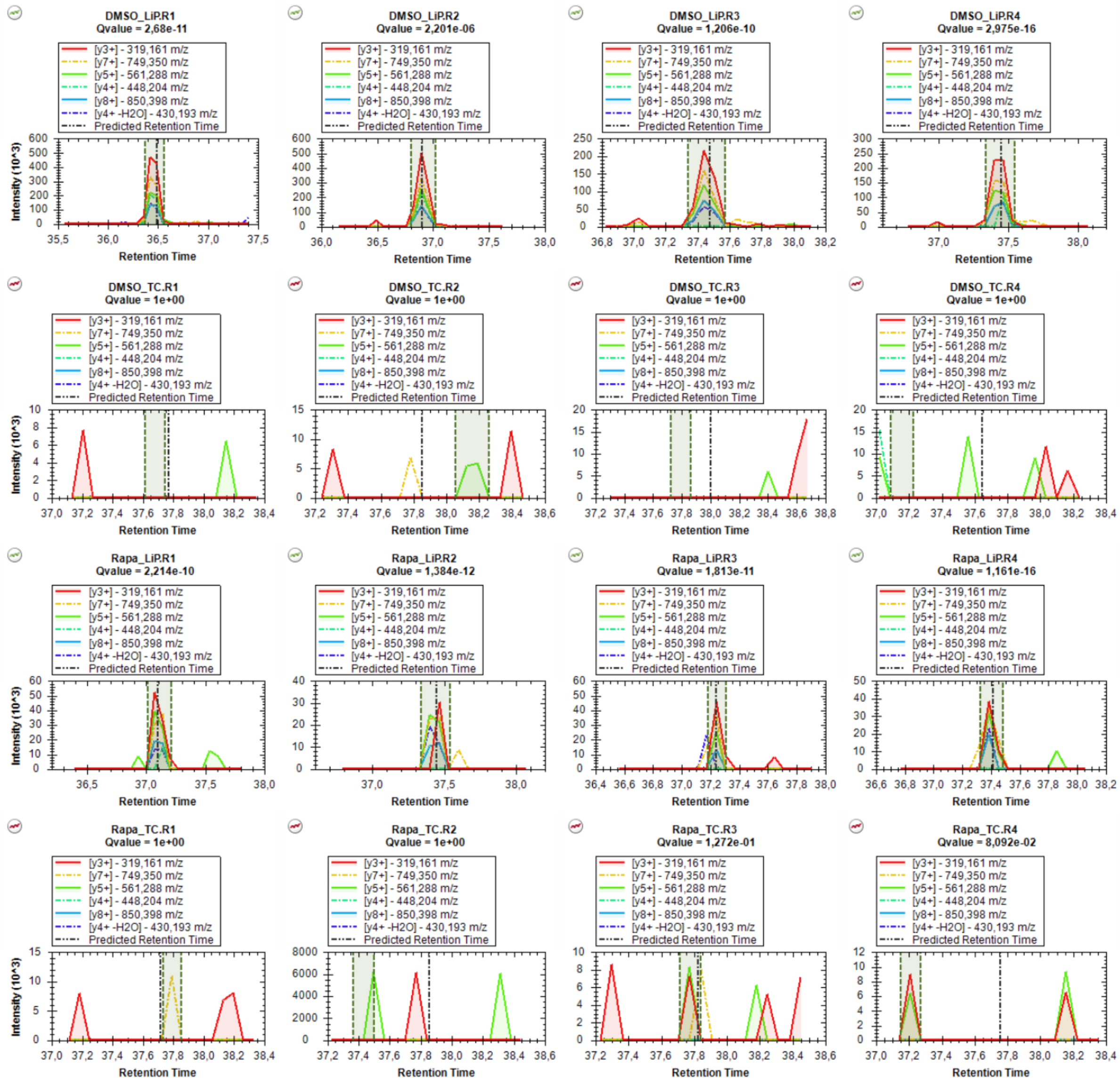


#### Annotated versus predicted spectrum

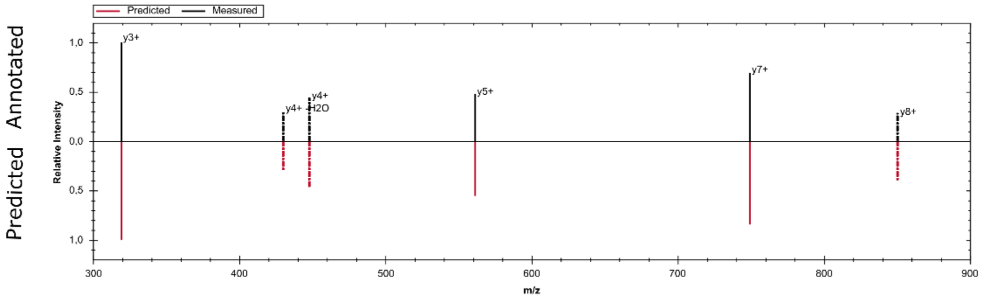

### QEVIRGWEEGVAQM2

Semi-tryptic; present in LiP conditions

DIA-NN v2.3, visualization in Skyline

**DMSO_LiP.R1**

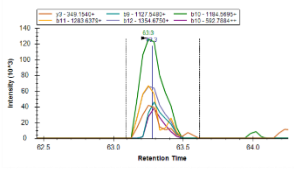

**DMSO_LiP.R2**

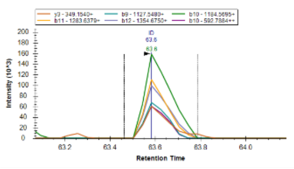

**DMSO_LiP.R3**

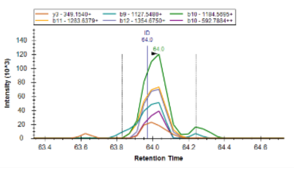

**DMSO_LiP.R4**

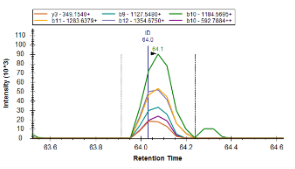

**DMSO_TC.R1**

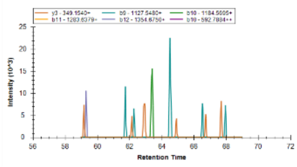

**DMSO_TC.R2**

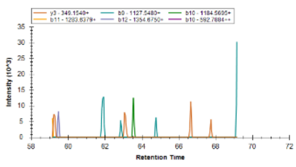

**DMSO_TC.R3**

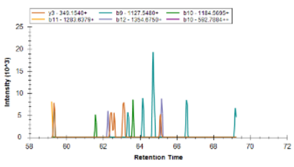

**DMSO_TC.R4**

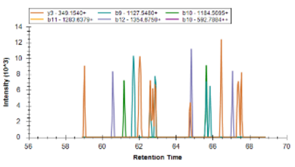

**Rapa_LiP.R1**

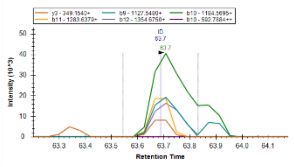

**Rapa_LiP.R2**

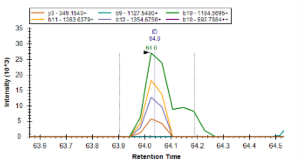

**Rapa_LiP.R3**

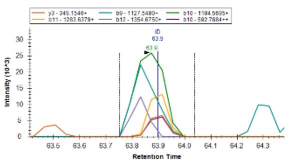

**Rapa_LiP.R4**

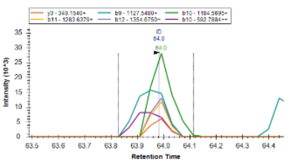

**Rapa_TC.R1**

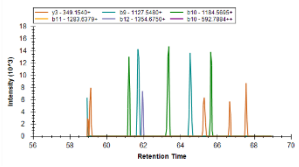

**Rapa_TC.R2**

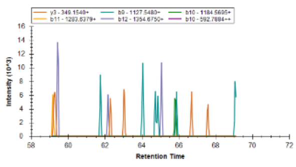

**Rapa_TC.R3**

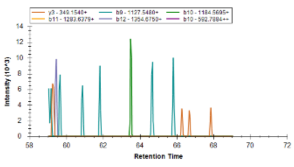

**Rapa_TC.R4**

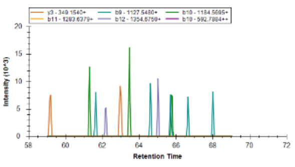

#### Annotated versus predicted spectrum

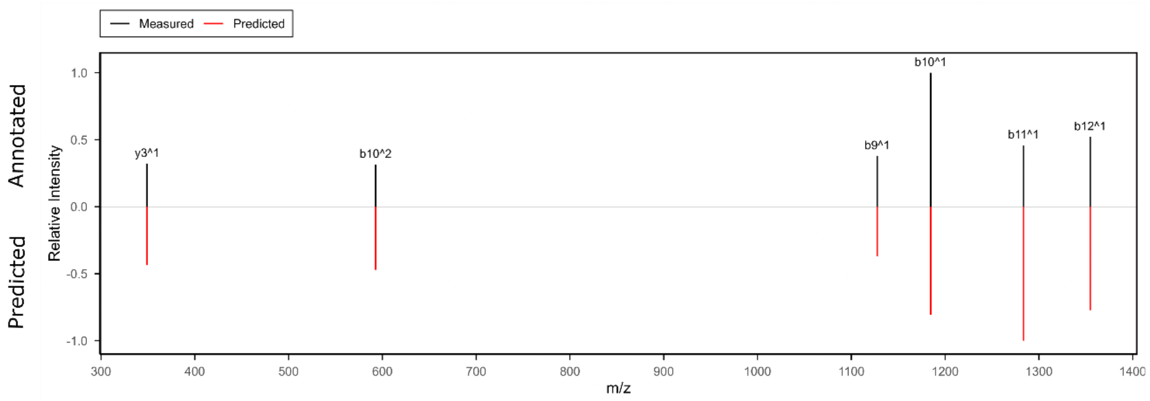

### QEVIRGWEEGVAQM2

Semi-tryptic; present in LiP conditions

Spectronaut v20.3

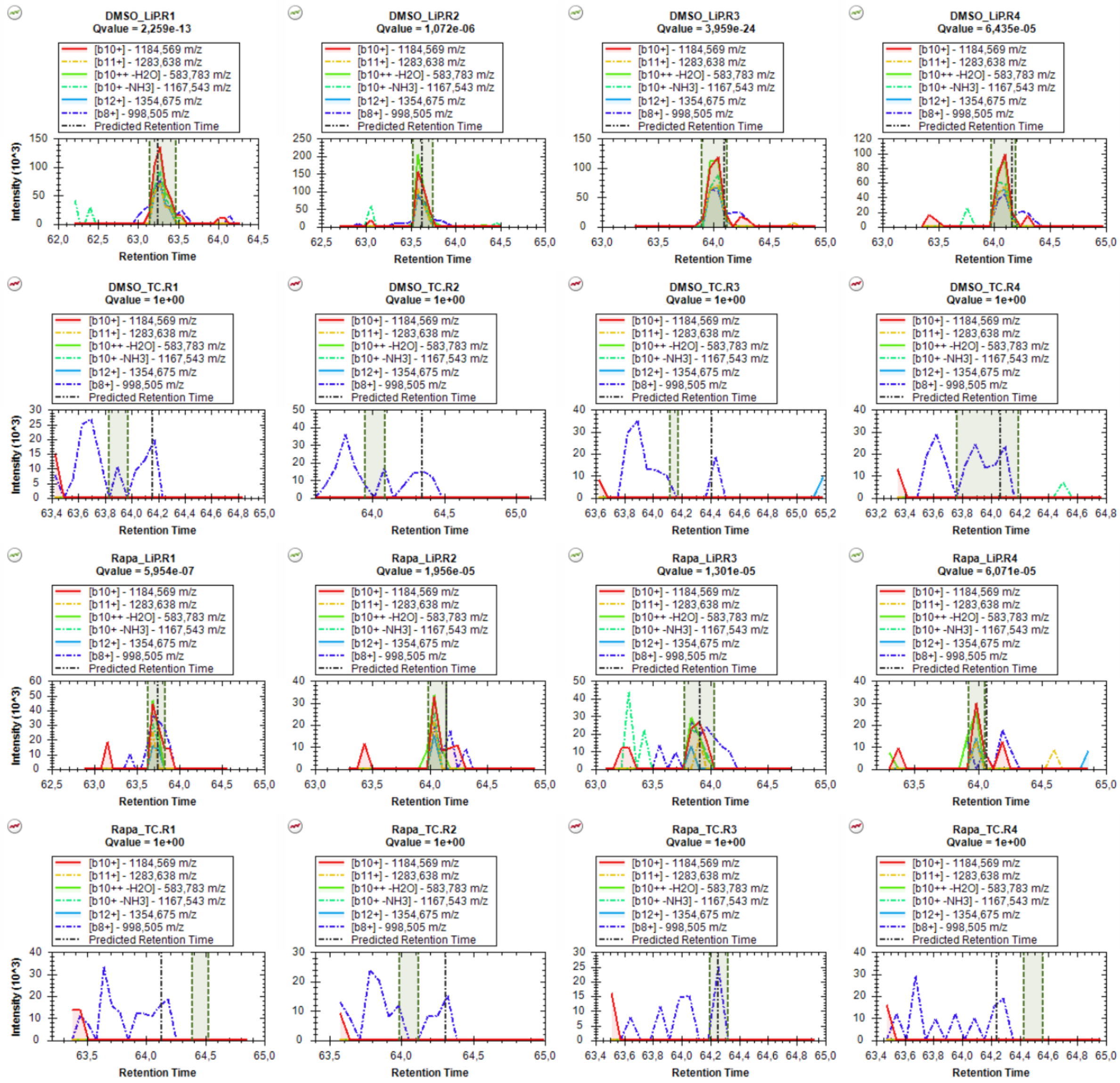


#### Annotated versus predicted spectrum

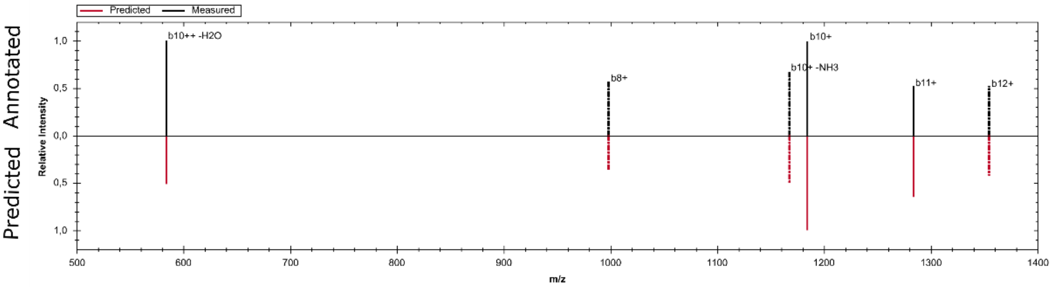

### QEVIRGWEEGVAQMSVGQR3

Tryptic; present in all conditions

DIA-NN v2.3, visualization in Skyline

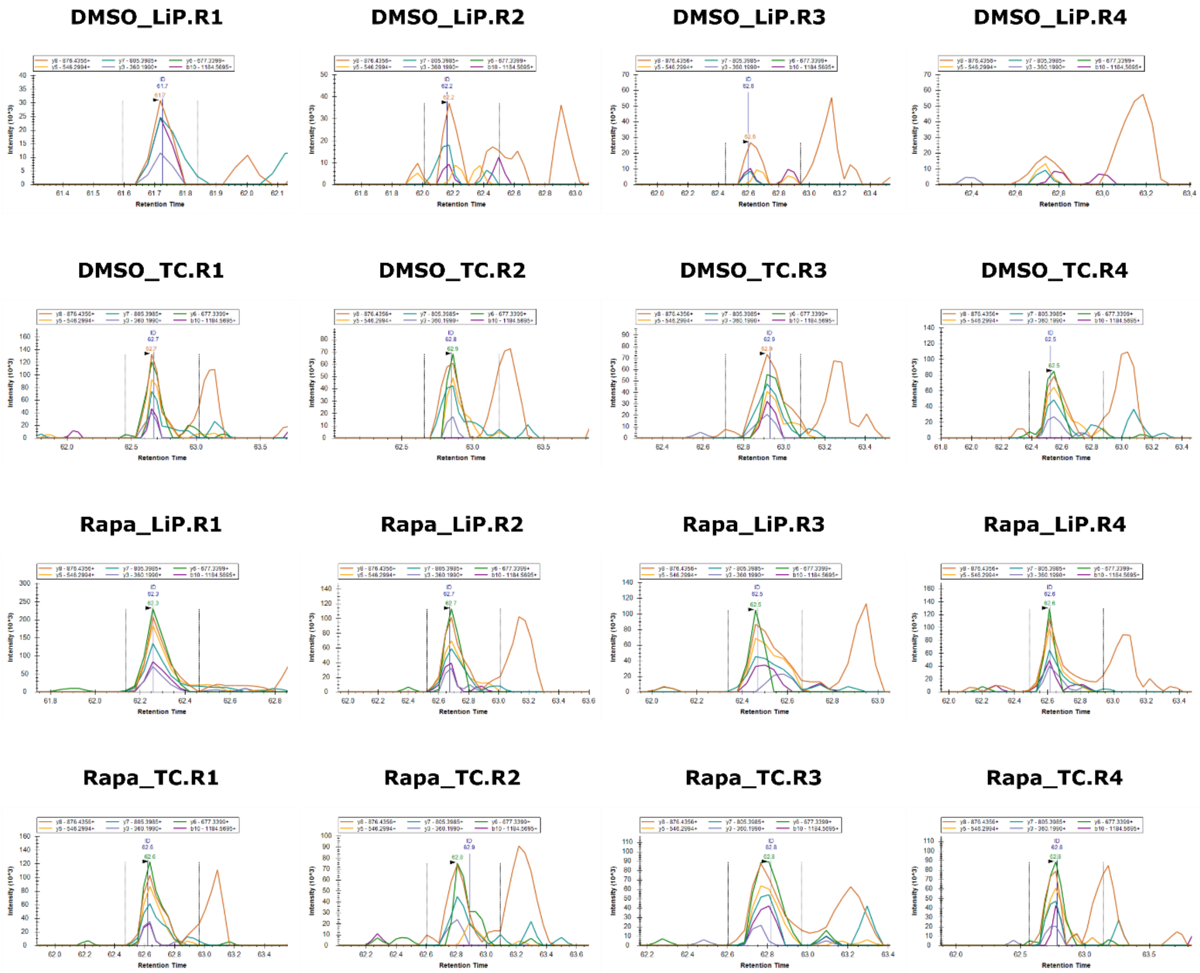


#### Annotated versus predicted spectrum

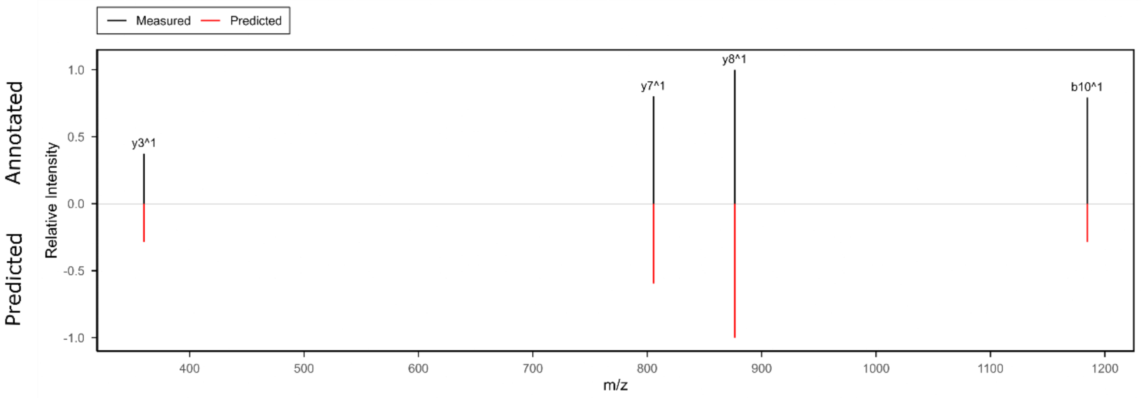

### QEVIRGWEEGVAQMSVGQR3

Tryptic; present in all conditions

Spectronaut v20.3

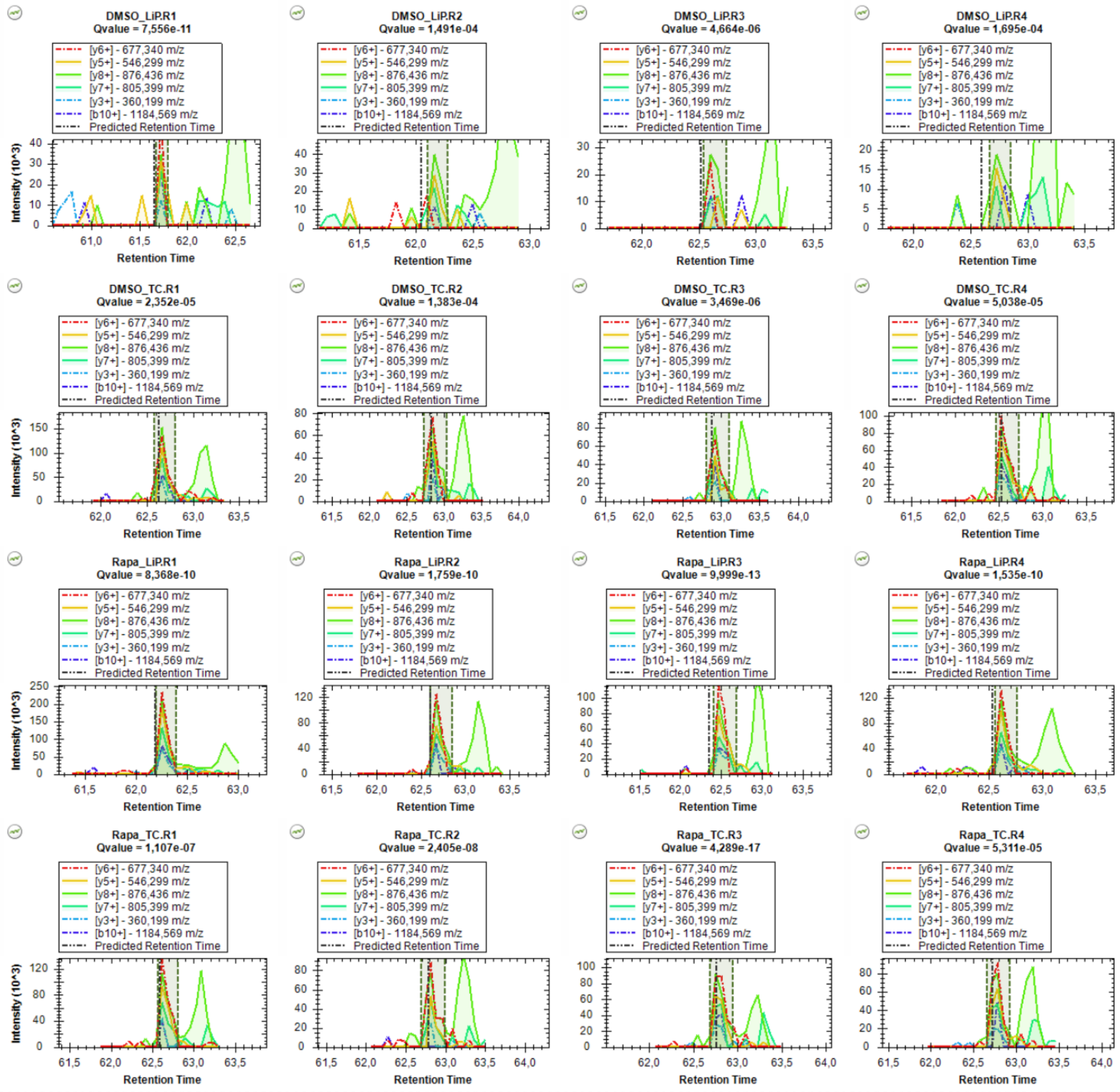


**Annotated versus predicted spectrum**

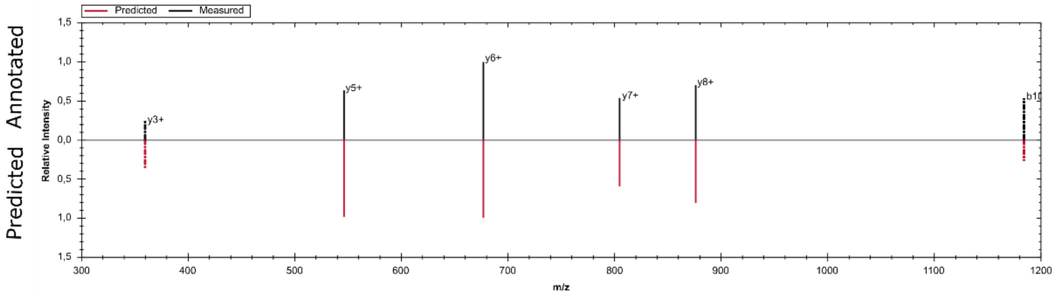

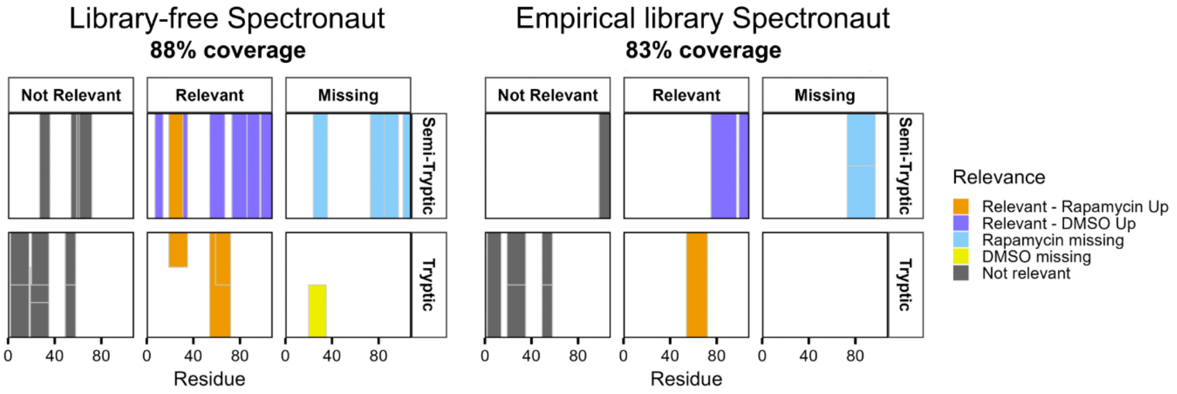


**Figure S5: Extended structural barcodes for FKBP1A in library-free and empirical Spectronaut workflows.** Each panel represents the FKBP1A sequence segmented by precursor trypticity (rows: semi-tryptic, tryptic) and detection status (columns: not relevant, relevant, missing). Relevance thresholds: $|\log_2(FC_{LiP}/FC_{TC})| \geq 1$ and adjusted p-value $\leq 0.05$. Bars indicate precursor positions along the protein sequence, coloured by relevance: orange (relevant and upregulated in rapamycin-treated LiP upon correction for TC), dark blue (relevant and upregulated in solvent control LiP upon correction for TC), grey (not relevant), light blue (missing in rapamycin-treated), yellow (missing in solvent control), white (not detected). Bars are stacked when there are multiple precursors corresponding to the same peptide sequence. Coverage percentages for each workflow are shown above the barcodes.

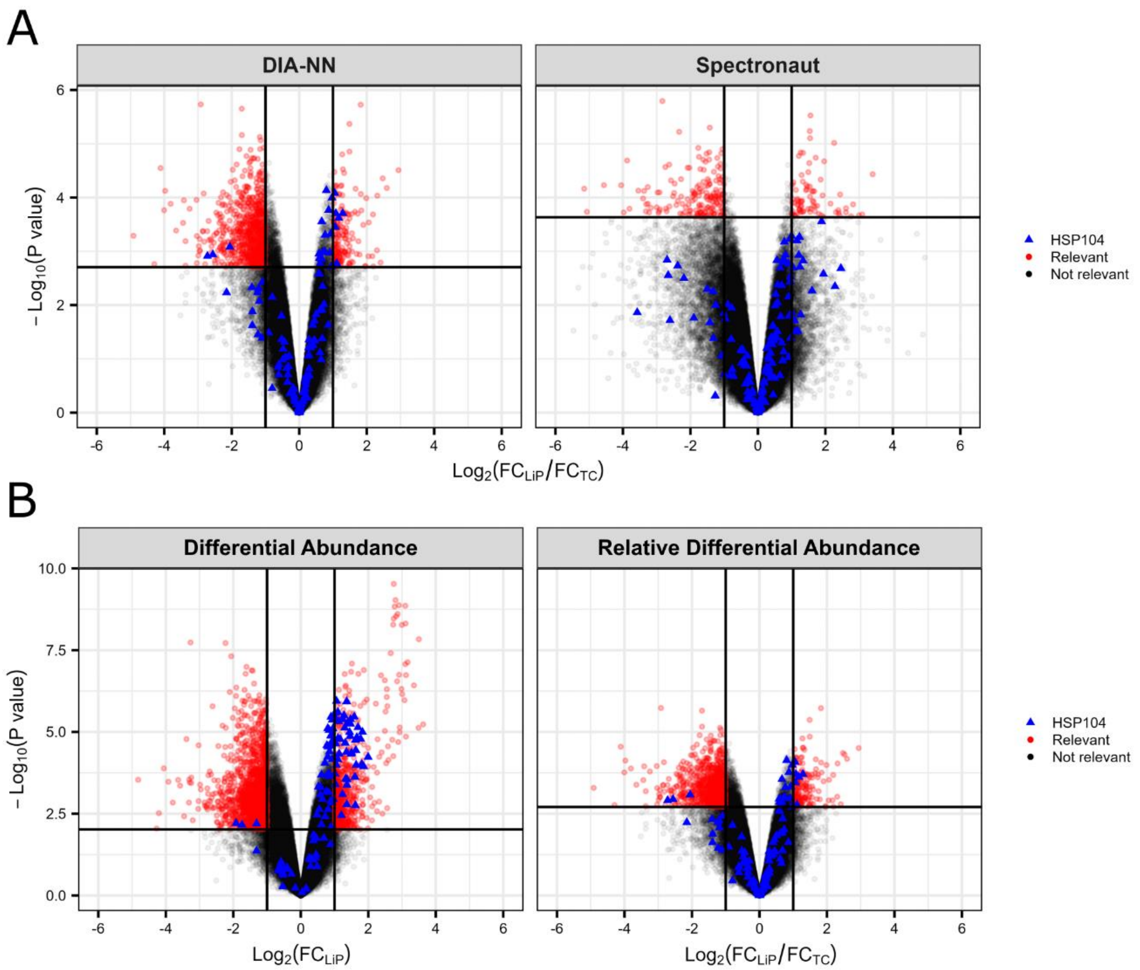


**Figure S6: Effect of search engine and relative abundance modeling in the yeast dataset. A)** Volcano plots comparing library-free DIA-NN and Spectronaut results (paired design). **B)** Volcano plots illustrating library-free DIA-NN results based on differential abundance (left) or relative differential abundance (right). In all panels, Hsp104 precursors are marked in blue, other relevant precursors in red, and non-relevant precursors in black. Relevance thresholds are indicated by horizontal and vertical lines: adjusted p-value ≤ 0.05 and $|\log_2(FC_{LiP}/FC_{TC})| \geq 1$ (RDA) or $|\log_2 FC| \geq 1$ (DA).

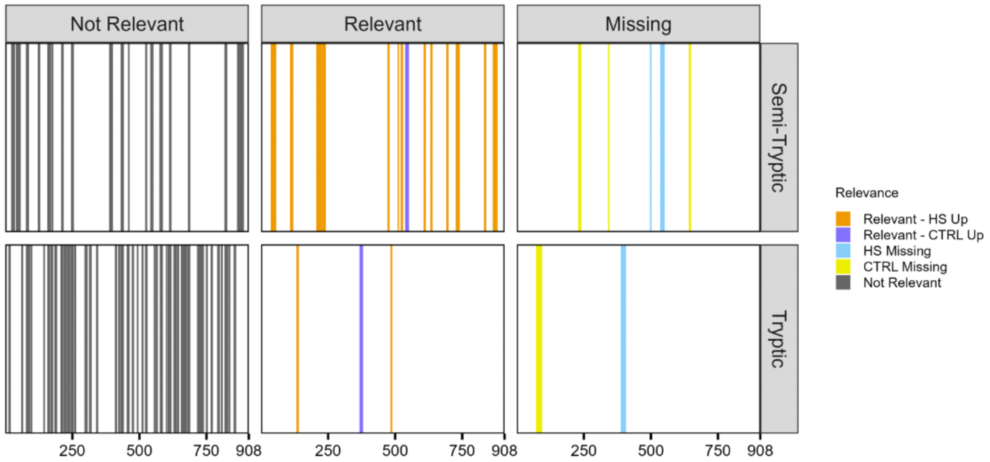


**Figure S7: Extended structural barcode for Hsp104 in library-free DIA-NN.** Each panel represents the Hsp104 sequence segmented by precursor trypticity (rows: semi-tryptic, tryptic) and detection status (columns: not relevant, relevant, missing). Relevance thresholds: $|\log_2(FC_{LiP}/FC_{TC})| \geq 1$ and adjusted p-value $\leq 0.05$. Bars indicate precursor positions along the protein sequence, coloured by relevance: orange (relevant and upregulated in rapamycin-treated LiP upon correction for TC), dark blue (relevant and upregulated in solvent control LiP upon correction for TC), grey (not relevant), light blue (missing in rapamycin-treated), yellow (missing in solvent control), white (not detected).

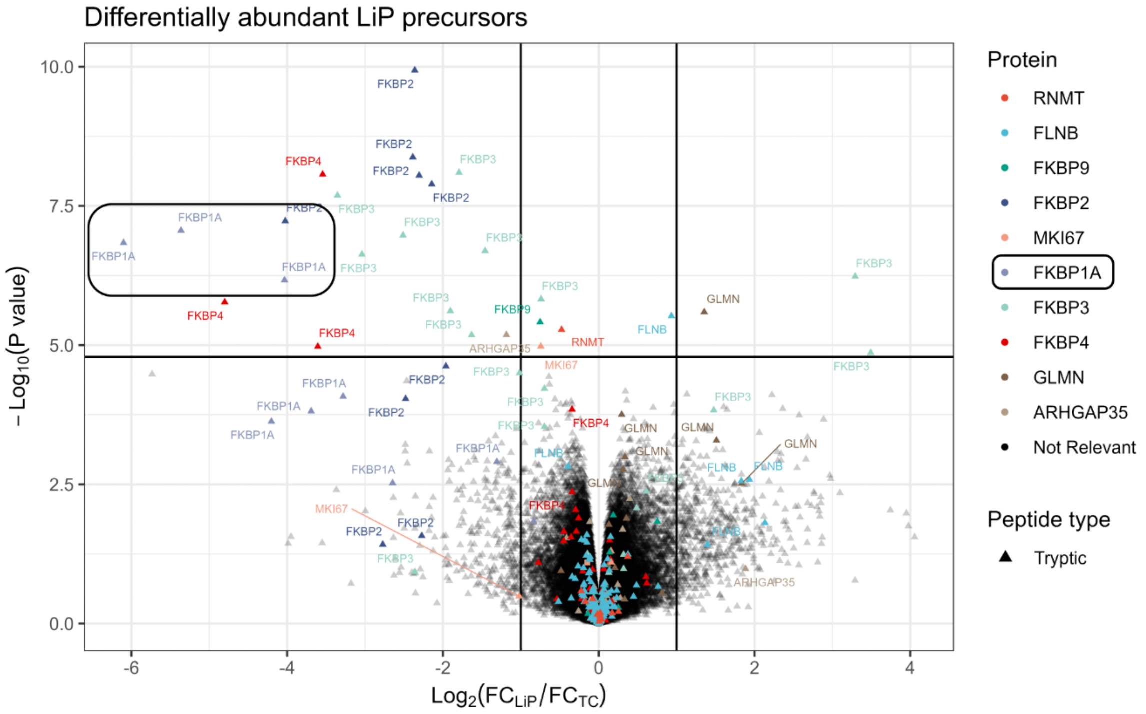


**Figure S8: DIA-LiPA analysis of the PELSA dataset (PXD034606).** Volcano plot showing the relative differential abundance between rapamycin-treated versus solvent (DMSO) control samples in the PELSA dataset, analysed using a minimally adapted version of DIA-LiPA. Relevance thresholds: $|\log_2 FC| \geq 1$ and adjusted p-value $\leq 0.05$. Each point represents a precursor, coloured by protein group. All detected peptides are fully tryptic, consistent with the PELSA workflow design, and no semi-tryptic peptides were identified. Because PELSA lacks trypsin-control samples and relies solely on long tryptic peptides, the analysis reduces to a standard differential abundance workflow rather than a structural LiP-MS readout.

**Table S1: Impact of HTRMS conversion and Semi-Specific Pipeline settings on semi-tryptic identifications in Spectronaut 20.3.**

| | **FULL ENUMERATION – HTRMS** | **FULL ENUMERATION – RAW** | **SMART ENUMERATION – HTRMS** | **SMART ENUMERATION – RAW** |
|---|---|---|---|---|
| ***Precursor IDs*** | 70K LiP<br>64K TC | 62K LiP<br>57K TC | 65K LiP<br>71K TC | 45K LiP<br>69K TC |
| ***Protein IDs*** | 5100 LiP<br>5400 TC | 4900 LiP<br>5100 TC | 5400 LiP<br>5900 TC | 5200 LiP<br>5800 TC |
| ***% semi-tryptics*** | **38% LiP**<br>5% TC | **35% LiP**<br>4% TC | **29% LiP**<br>3% TC | **5% LiP**<br>1% TC |
